# Human disturbances erode the diversity of species resilience strategies

**DOI:** 10.1101/2021.09.29.462372

**Authors:** Thomas Merrien, Katrina J Davis, Moreno Di Marco, Pol Capdevila, Roberto Salguero-Gómez

## Abstract

Human activities are drastically reshaping Earth’s ecosystems. Across the tree of life, species become threatened and ultimately go extinct when they are unable to cope with these changes. Hence, understanding the resilience of natural populations is necessary to understand and predict species’ capacity to cope with increasing human pressures. Here, we use high-resolution demographic information for 921 populations of wild plants and animals to assess how they respond to increasing levels of human pressure. We show that fewer successful resilience strategies, allowing population to persist in disturb environments, exist in human-influenced habitats compared to pristine habitats. In contrast, pristine habitats host species with higher resistance and faster recovery than more altered environments. Importantly, the examined macroecologial patterns of demographic resilience are kingdom- and mobility-specific: natural populations of plants recover faster and have a propensity to grow faster after a disturbance (*i.e*., compensation) in urban areas than in pristine habitats, while these tendencies do not appear in animals. Likewise, populations of animals with limited mobility are less able to resist or compensate for disturbances in human altered environments than highly mobile populations. Our results suggest that human activities have eroded the diversity of natural populations’ resilience strategies. This finding implies that species will be less tolerant to disturbance in the future, as continuing biodiversity loss and increasing human impacts will ultimately shrink the spectrum of resilience strategies of organisms.

---

The exponential growth that has characterised human populations since the industrial revolution has fundamentally modified Earth’s life support systems^1,2^. Examples of such human impacts include rapid increases in agricultural areas, now accounting for 37% of the land surface^3^, or the growth of human-made objects (buildings, infrastructure, products, etc.), which now exceeds the cumulative mass of all living organisms worldwide^4^. Recent estimates suggest that 59% of the Earth’s terrestrial surface is negatively impacted by human land use^5^, leading to rapid decreases in species’ ranges^6^. For example, by 2100 the distribution of black spruce (*Abies nigra*) in France is projected to shrink by 80%^7^, and the habitat of the Snow leopard (*Panthera unica*) in the Himalayas to decline by 30%^8^. More broadly, land-use change is responsible for the loss of 5-16% of vertebrate species compared to pristine habitats^9^, and has contributed to a 60% increase in vascular plant extinctions between 1900 and 2015^10^. Ultimately, these fast and large-scale changes in biodiversity affect the dynamics of Earth’s ecological systems^11^ and their services^12^. In fact, the homogenisation of the environment, caused by land-use modification, may result in the loss of functional diversity^13^.

The impacts of human land use on the viability of natural populations can be directly quantified by assessing changes in the demography of species. Changes in a natural population’s vital rates (survival, development, reproduction) shape its ability to persist in a given location, ultimately determining a species’ overall spatial distribution^14^. As such, demographic approaches are powerful tools to examine population responses to human disturbances^15^. Indeed, extensive evidence exists that disturbances shape species’ life-history strategies (*i.e*., schedules of vital rates along a species’ life cycle^16^), and that these strategies in turn determine the viability of natural populations^17^. Furthermore, in recent decades, life-history theory has developed frameworks to quantify and classify species’ responses to disturbances using organismal traits^18,19^. Traits such as generation time or age at maturity impact population performance^19^. However, identifying the life-history strategies that enable species to resist and recover from disturbances has become a promising research focus in better understanding the resilience of natural populations to human impacts^20^. Some studies highlight the key role of life-history traits, such as the mean reproductive output or generation time, in determining the demographic resilience of species^20^. However, it still remains unclear how do human activities alter life history strategies and its consequences for the demographic resilience of species.

Here, we examine how the resilience of 921 natural populations of 279 plants, 1 alga, 45 animals, and 1 fungal species worldwide correlates with human activities. To do so, we build a spatially and phylogenetically explicit model parameterised with life-history traits and transient (*i.e*., short-term) dynamics metrics that describe how populations inherently respond to disturbances^21^ (hereby *demographic resilience*). This demographic information was obtained from the open-access COMPADRE^22^ and COMADRE^23^ databases, coupled with high-resolution human impact information from the Human Footprint database^24^. We hypothesise that: (H1) species’ populations located near urban areas have a greater ability to resist, recover, and even increase in size directly after the disturbance (*i.e*., compensate) compared to populations in more pristine habitats. These three aspects: resistance, speed of recovery, and compensation being the key defining characteristics of demographic resilience^20^. This greater demographic resilience would have evolved via life history trait changes to allow these populations to persist in heavily disturbed, human habitats^25^; (H2) the degree to which human impacts have shaped a species’ demographic resilience depends on each species’ capacity for dispersal. Specifically, we expect the impact of human activities on demographic resilience to be stronger in animals with limited mobility and all plants since human disturbances are inescapable to its established individuals; and (H3) human pressures constrain the repertoire of life-history strategies that confer high demographic resilience to fewer viable combinations. Indeed, in general, human activities homogenise habitats (*e.g*., via urban development and agricultural monocultures^26^), and hence these activities might lead to a convergence in demographic responses to disturbances.

## The resilience hypervolume: human activities constrain the spectrum of resilience strategies

To assess the main human impacts on terrestrial habitats to species’ life-history traits and their demographic resilience, we reduced the dimensionality of the eight human activities quantified by the Human Footprint^24^ index using a Principal Component Analysis (PCA). We retained two dominant axes of variation of human activities following the Kaiser criterion^27^ (*i.e*., associated eigenvalues >1, Table S5), corresponding to human presence and agricultural land use. These two axes together explain >52% of the variance in human activities (Fig. 1). The first axis (PC1), which absorbs 35.86% of this variance, describes human presence, separating areas according to human population density, degree of built environment, and light pollution. The second axis (PC2) absorbs 16.63% of the variance and describes land use, separating extensive pastures from intensive agricultural croplands.

**Figure 1.**
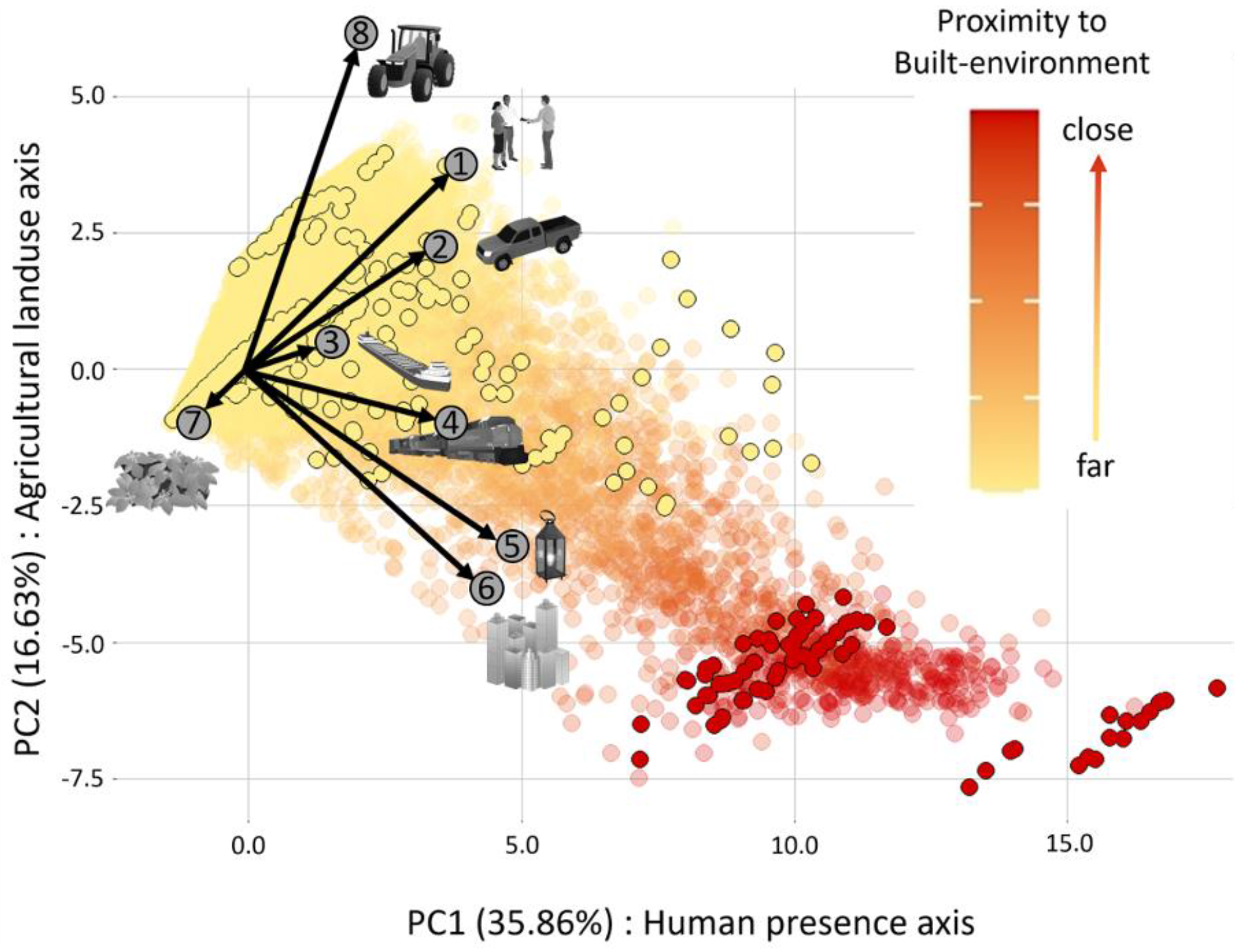
Principal Component Analysis (PCA) plot of terrestrial habitats worldwide, derived from the Human Footprint^33^. The PCA shows the two dominant axes of variation: PC1 corresponds to variation on human presence, and it is positively correlated with (1) human population density, (2) roads, (3) navigable waterways, (4) railways, and (5) light pollutions, (6) built environment, and it explains 35.86% of variation. PC2 differentiates locations according to land use, and it represents a trade-off between (7) extensive pastures, and (8) intense croplands, and it explains 16.63% of variation. Each dot corresponds to a terrestrial unit of 10 km^2^ and is colour-coded according to its distance to a human-built environment, here defined as human produced areas that provide the setting for human activity (delimited using satellite imagery)^33^. Dots with a black circle correspond to our examined 921 natural populations.

When considered together, the demographic resilience of the examined 921 natural populations is strongly shaped by human presence and agricultural land use. To estimate species’ demographic resilience, we used a framework we recently developed^21^ based on well-established metrics of transient (*i.e*., short-term) dynamics^28^. For any species, this framework quantifies a population’s ability to resist, recover, or even increase in size when affected by a disturbance (compensation) (See Extended Methods). Next, we examined how resistance, recovery, and compensation change along the axes of human pressure to test whether species located near highly disturbed habitats are more resistant, recover faster or compensate more (H1). Along the human presence axis (PC1), populations that are closer to human settlements are significantly more resistant (linear model: *t-test*_df_=913 = 2.098, *P* = 0.036, Fig. 4a), while their compensation and speed of recovery are independent of PC1 (compensation: *t*_913_ = -1.342, *P* = 0.180; recovery: *t*_913_ = 1.597, *P* = 0.111) (Fig. 4; Table S6). With regards to agricultural land use, species located closer to intensive agricultural areas have higher compensation than those in pristine or extensive areas (t_913_ = 2.767, P = 0.006, Table S6), but their resistance and speed of recovery are on average independent of land use (PC2).

**Figure 2.**
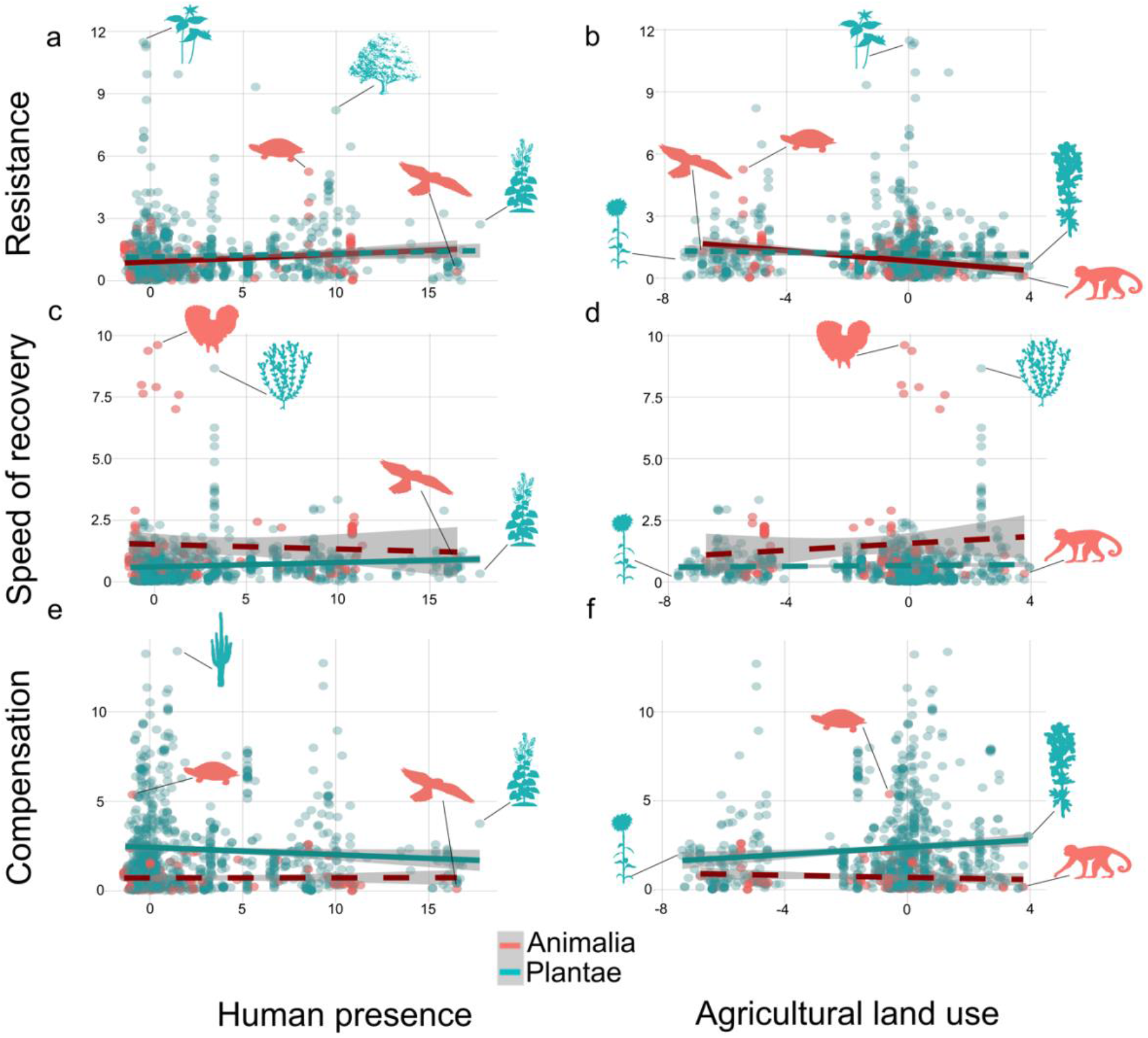
The three components of demographic resilience –resistance, recovery, and compensation– are affected by human pressure in different ways for plants than for animals. Correlation of three key demographic resilience components of our studied 921 natural populations along the continua of human presence (a, c, e) and agricultural land use (b, d, f) (See Figure 1). The shown linear fits describe the general correlations (orange: animals; green: plants), while the grey area represents the 95% confidence intervals. A selected number of species are represented by silhouettes, from left to right: (a) *Trillium persistens* (persistent trillium), *Emydura macquarii* (Macquarie turtle), *Cornus florida* (flowering dogwood), *Falco naumanni* (lesser kestrel), *Alliaria petiolata* (garlic mustard); (b) *Echinacea angustifolia* (narrow-leaved purple coneflower), *Falco naumanni, Emydura macquarii, Trillium persistens, Dracocephalum austriacum* (Austrian dragon’s head), *Cebus capucinus* (Colombian white-face capuchin); (c) *Bonasa umbellus* (ruffed grouse), *Arenaria serpyllifolia* (thyme-leaf sandwort), *Falco naumanni*, *Alliaria petiolata*; (d) *Echinacea angustifolia, Falco naumanni, Bonasa umbellus, Arenaria serpyllifolia*, *Dracocephalum austriacum, Cebus capucinus*; (e) *Podocnemis expansa* (Arrau turtle), *Pseudomitrocereus fulviceps, Falco naumanni* ; (f) *Echinacea angustifolia, Podocnemis expansa, Dracocephalum austriacum, Cebus capucinus*.

**Figure 3.**
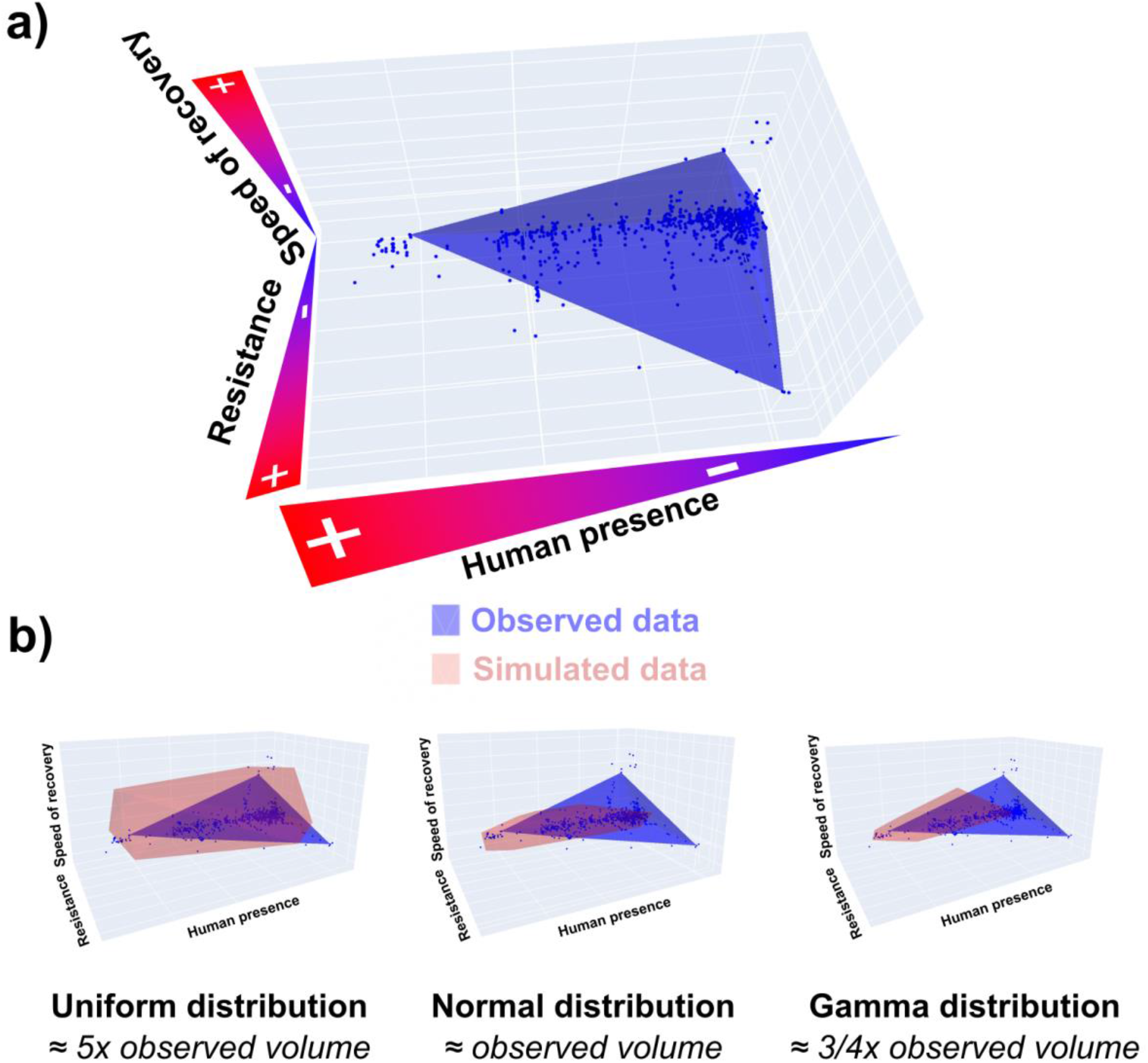
Human presence tightened the repertoire of resilience strategies. a) Plot of the 3D framework composed of human presence, recovery time, and resistance (compensation is not displayed here for visual convenience). b) Comparisons of the volume of achieved strategies in the space defined by compensation – resistance – recovery time – human presence under the hypothesis of a normal distribution of the resistance and speed of recovery, a uniform distribution of the three same traits and the observed distributions in our populations (for graphical reason only 3D framework without compensation are displayed).

**Figure 4.**
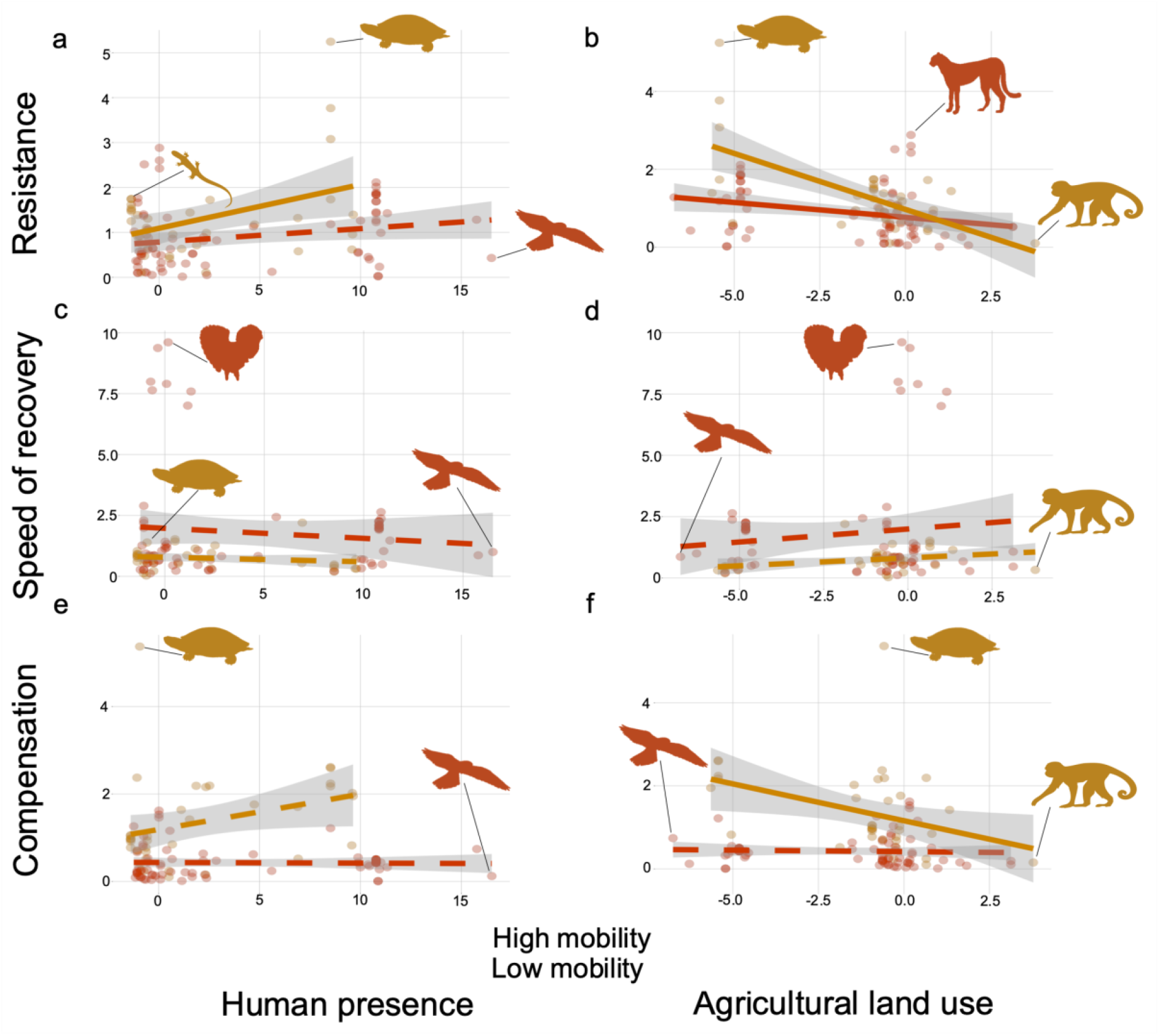
low mobility/sessile species’ resilience is more strongly affected by human pressure. Correlation of three key demographic resilience components (resistance, speed of recovery, and compensation) of our studied 107 natural populations along the continua of human presence (a, c, e) and agricultural land use (b, d, f) (See Figure 1). The shown linear fit describes the general correlations (brown: high mobility [>100 km]; yellow: low mobility/sessile), with the grey area representing 95% confidence intervals. A selected number of species are represented by silhouettes, from left to right: (a) *Sceloporus arenicolus* (dune sagebrush lizard), *Emydura macquarii, Falco naumanni*; (b) *Emydura macquarii, Acinonyx jubatus* (cheetah), *Cebus capucinus*; (c) *Podocnemis expansa, Bonasa umbellus*, *Falco naumanni*; (d) *Falco naumanni, Bonasa umbellus, Cebus capucinus*; (e) *Podocnemis expansa, Falco naumanni* f) *Falco naumanni, Podocnemis expansa, Cebus capucinus*.

The impacts of human activities on demographic resilience differ considerably between plants and animals. On average, animal populations close to human presence tend to be more resistant than plant populations (*t*_107_ = 2.357, *P* < 0.010; phylogenetically and spatially corrected models: *P_MCMC_* = 0.032; Fig. 2a; Table S7; Table S11), but exhibit lower resistance in intense agricultural areas (*t*_107_ = -3.872, *P* < 0.001 ; Fig. 2b; Table S7), while their speed of recovery is not affected by human presence (t_107_ = -0.526, P = 0.600) nor agricultural land use (t_107_ = 0.917 and P = 0.343; Fig 2.c&d, Table S7). The compensatory potential of animals is also unaffected by human presence (t_107_ = 0.064, P = 0.949; Fig. 2e) and agricultural land use (t_107_ = -0.952, P = 0.343; Fig. 2f, Table S7). In contrast, while the resistance of plant populations is insensitive to human presence (*t*_788_ = 1.508, *P* > 0.050; Fig. 2a) and agricultural land use (*t*_788_ = -0.889, P > 0.050, Fig. 2b ; Table S7), their recovery post disturbance is faster (*t*_788_ = 2.976, *P* < 0.010 ; Fig. 2c), and has lower compensatory potential closer to human presence (*t*_788_ = -2.07, P = 0.039, *P_MCMC_* < 0.010; Fig. 2e). However, plants’ compensation and speed of recovery are not affected by agricultural land use (Fig. 2f).

The repertoire of resilience strategies defined by the three components of demographic resilience –resistance, compensation, and speed of recovery– drastically decreases in more densely human populated areas (Fig. 3a). Indeed, the variance in resilience strategies decreases in areas with more prominent human activities. To test whether this trend is statistically supported, we compared and contrasted different null models using convex hulls^29^ (Fig. 3b). This approach assumes different potential distributions regarding how resistance, speed of recovery, and compensation co-vary with human presence axis (PC1 in Fig. 1). The observed repertoire of resilience strategies is significantly different from that expected under a normal, gamma, or uniform distribution that corresponds to a hypothetic world where species with high and low values of demographic resilience would exist in all kinds of areas, including areas with a high human pressure (Fig. 3b; See also https://colab.research.google.com/github/merrienthomas/HFP_appendix/blob/main/display_fig.ipynb for an interactive shiny app representation of the 4-D space). Indeed, the volume occupied by the resistance, speed of recovery, and compensation of our 921 natural populations as a function of their proximity to human settlements under a uniform distribution is five-fold larger than the volume occupied by our data (Monte Carlo test *P* = 0.001), 0.75-fold smaller under a gamma distribution (P = 0.001), but similar under a normal distribution (*P* = 0.348). However, despite having a similar volume, the volume that would emerge from a set of normal distributions does not fully overlap with our observed data (overlap is 77%). The distribution of the three components of demographic resilience along agricultural land use behaves similarly to the distribution along human presence with a thinning (decreased volume) of resilience diversity in areas with intensive agricultural practices. One exception is that both the normal and gamma distributed models have volumes equivalent to the volume of the observed data (both P > 0.050), but the percentage of overlap is close to 87% for both of them (https://colab.research.google.com/github/merrienthomas/HFP_appendix/blob/main/display_fig.ipynb). Concerning the normal and gamma models, they fail to represent the diversity of strategies in pristine habitats while they overestimate diversity in highly anthropized habitats. Although the gamma model better captures the shape of the observed data, it overestimates the overall volume of its convex hull. This finding suggests that our estimates of demographic resilience are not a random representation of all possible strategies, and the resilience repertoire of the examined 921 natural populations has been eroded by human activities.

## Mobility to escape from human disturbances

The ability of species to move long-distances is an important factor in determining their demographic resilience in our study. To test (H2) whether species with the ability to flee from a disturbance may require lower levels of demographic resilience compared to those with limited mobility, we re-examined the data set with our spatially and phylogenetically explicit models (results provided in terms of P_MCMC_, as these methods use a Bayesian framework). We also used classic linear regression (classical P-values), this time comparing animal species with long-distance moving ability (>100 km; e.g. *Bonasa umbellus*, ruffed grouse; *Falco naumanni*, lesser kestrel) *vs*. those without it (<100 km; *Emydura macquarii*, Macquarie turtle; *Sceloporus arenicolus*dune sagebrush lizard). Both short-mobility/sessile and long-mobility species are or tend to be more resistant when they are closer to human presence (*t*_34_ = 2.510, *P* = 0.017; *t*_71_ = 1.954, *P* = 0.055; respectively; Table S7, Fig 4a). However, as expected, the increase in resistance is more pronounced for less mobile species than for more mobile ones (Fig. 4a). Along the axis of agricultural land use, both groups show a sharp decrease in resistance in intense agricultural lands (*t*_34_ = -4.964, *P* < 0.001 for low mobility/sessile; *t*_71_ = -2.362, *P* = 0.021 for high mobility; Table S7, Fig4b). Although fewer mobile species are more resistant than high mobility species to changes in agricultural land use (*P_MCMC_* = 0.04, Table S11). The speed of recovery and compensation of low mobility/sessile species tends to change in response to the axes of human activity (Fig. 4c,d,e&f). Specifically, in these species, the speed of recovery shows a tendency to increase with agricultural land use intensity (*t*_34_ = 1.916, *P* = 0.064; Table S7, Fig. 4d), and their potential for compensation tends to increase closer to human presence (*t*_34_ = 1.971, *P* = 0.057; Table S7, Fig. 4e) while compensation decreases closer to agricultural areas (*t*_34_ = -2.374, *P* = 0.023; Table S7, Fig. 4f). Finally, as hypothesised, the speed of recovery and compensation in high mobility species is not affected by either axis of human pressure (Fig. 4c,d,e&f).

## Impacts of Human pressure on life-history traits

To test (H3) whether human pressures correlate with changes in underlying life-history traits that are known to control the three components of demographic resilience^20,30^ (See Methods and Supplementary Materials), we used spatially and phylogenetically explicit MCMC generalized linear mixed regression models. The generation time of species with high mobility is significantly longer in populations close to human presence and high-intensity agricultural land (*P_MCMC_* < 0.050; Table S11), while it remains unaffected in less mobile species (Table S6 & S7). This shows that longer generation times can be favoured in highly disturbed habitats when species have the possibility to move across long distances. However, for less mobile animal species, age at maturity (and consequently generation time) decreases significantly when their populations are located close to human presence and intensive agriculture (*P_MCMC_* < 0.050). Reaching maturity at a younger age would allow individuals to better deal with habitat disturbances by producing offspring earlier. Plant populations closer to human-transformed habitats tend to have higher rates of individual-level shrinkage than those in pristine habitats (Table 1a), implying that shrinkage might be a strategy to better cope with habitats disturbed by humans.

**Table 1.** The effects of human pressure are restricted to only a few demographic processes and life-history traits. Summary of the results of the trait analysis for plants (light blue) and animals (light red) using the residual method (i.e., the spatially and phylogenetically explicit method) described in Methods. The human presence axis and agricultural land use axis refer to the two main dimensions of human activities from the Human Footprint database^33^, as detailed in Figure 1. Result are given in these two columns according to the following format: the effect size (*P_MCMC_*-values). The columns “size correction” and “spatial correction” indicate whether we corrected for adult body size and spatial autocorrelation in the model (✓ and ✗ symbols respectively stands for when corrections are included and not included) (See Materials and Methods for further details on model choice). Note that none of the examined animal populations include shrinkage or clonal reproduction (thus “-“). *P_MCMC_*-values parentheses with underline: *P_MCMC_*<0.

| Residual method for phylogenetically and spatially corrected model for plants and animals |  |  |  |  |
| --- | --- | --- | --- | --- |
| Trait | Human presence axis | Agricultural land use | Size correction | Spatial correction |
| a) Vital rates |  |  |  |  |
| Survival | 0.003 (<0.05) | -0.001 (>0.05) | ✗ | ✓ |
|  | -0.010 (>0.05) | -0.029 (>0.05) | ✓ | ✗ |
| Growth | -0.001 (>0.05) | 0.002 (>0.05) | ✓ | ✓ |
|  | 0.004 (>0.05) | 0.009 (>0.05) | ✓ | ✓ |
| Shrinkage | 0.013 (>0.05) | 0.006 (>0.05) | ✓ | ✓ |
|  | - | - | - | - |
| Sexual reproduction | -0.038 (>0.05) | 0.044 (>0.05) | ✓ | ✓ |
|  | -0.015 (>0.05) | -0.001 (>0.05) | ✓ | ✗ |
| Clonal reproduction | -0.008 (>0.05) | -0.017 (>0.05) | ✗ | ✓ |
|  | - | - | - | - |
| Shrinkage of pre-reproductive stages | 0.012 (<0.05) | 0.014 (>0.05) | ✗ | ✓ |
|  | - | - | - | - |
| Survival of reproductive stages | 0.001 (>0.05) | 0.002 (>0.05) | ✗ | ✓ |
|  | 0.005 (>0.05) | -0.007 (>0.05) | ✓ | ✓ |
| Growth of reproductive stages | 0.006 (>0.05) | 0.011 (>0.05) | ✓ | ✓ |
|  | 0.004 (>0.05) | -0.027 (>0.05) | ✓ | ✓ |
| b) Life-history traits |  |  |  |  |
| R<sub>0</sub>: Net reproductive rate | 0.034 (>0.05) | 0.042 (>0.05) | ✓ | ✓ |
|  | 0.057 (>0.05) | 0.126 (>0.05) | ✗ | ✗ |
| S: Degree of iteroparity | -0.036 (>0.05)<br>-0.001 (>0.05) | -0.047 (>0.05)<br>0.008 (>0.05) | X<br>✓ | ✓<br>X |
| T: Generation time | -0.001 (>0.05)<br>0.283 (<0.10) | -0.011 (>0.05)<br>0.509 (>0.05) | ✓<br>✓ | ✓<br>X |
| L<sub>α</sub>: Age at maturity | -0.023 (>0.05)<br>0.013 (>0.05) | -0.009 (>0.05)<br>-0.016 (>0.05) | ✓<br>✓ | ✓<br>✓ |
| H: Rate of actuarial senescence | 0.004 (>0.05)<br>-0.012 (>0.05) | 0.009 (>0.05)<br>-0.008 (>0.05) | ✓<br>✓ | ✓<br>X |
| p<sub>Rep</sub> : Probability of achieving reproduction before dying | 0.005 (>0.05)<br>0.002 (>0.05) | -0.002 (>0.05)<br>0.022 (>0.05) | ✓<br>X | ✓<br>✓ |

## Discussion

Our examination of the demographic resilience of plants and animal populations worldwide suggests an erosion in the diversity of resilience strategies in environments heavily disturbed by humans. Here, using high-resolution demographic data from 921 natural populations from 334 animal and plant species, we find that (1) urban areas host a limited diversity of demographically resilient strategies, with fewer populations with high levels of resistance or fast-recovering abilities located near human activities compared to populations in pristine habitats; (2) plant species are recovering faster but experiencing a lower compensation close to urban areas than in pristine environment, while for animals, these two aspects of resilience are not affected by human pressure; (3) the demographic resilience of species with limited mobility is more strongly affected by human presence than that of highly mobile species; and (4) the erosion of demographic resilience caused by human pressure is likely linked to their underlying life-history strategies: individuals of populations located closer to human settlements tend to decrease in size more frequently (*i.e*., shrink) and live faster (*i.e*., reproduce earlier) than those in pristine habitats.

Our study clearly shows that human pressure affects the ability of natural populations to cope with disturbances. Land-use change is a strong predictor of species’ evolution^31^, geographic range^32^, as well as extinction rates in mammals^33^ and plants^34^. Added to this growing body of literature evaluating novel dynamics in the Anthropocene, we find that human activities can also be a strong predictor of resilience strategies. Indeed, the correlates of human activities with different dimensions of population resilience —resistance, recovery, and compensation— show different directionality between plants and animals, and even between animals depending on their degree of mobility. However, our analyses confirm that human pressure, especially human presence, and agricultural land use, have already had a significant impact on species’ capacity to respond to disturbances. Ultimately, our analyses support recent evidence for the human-induced biotic homogenisation of natural systems^13^, here including resilience strategies. Our findings illustrate the pervasive impacts of human pressure on biodiversity, which spans multiple levels of biological organisation, from the demographic decline of individual populations to the loss of resilience strategies across different kingdoms^13,26^. Thus, some populations are able to cope with human changes and even to adapt, while for others, the speed of these changes can outpace their rate of adaptation^35^. This is intuitive as adaptation usually requires several generations and repetitive events of natural selection to occur^36^. For instance, evidence of phenotypic adaptation to urban environments has nevertheless already been shown in birds^1537^. Our results also support recent findings that species closer to human activities are more generalist than those in remote areas^38^. These impacts at macroecological scale showcase the long-term consequences of the Anthropocene for species ability to cope with disturbances.

Species with the most extreme values of demographic resilience in our study, surprisingly, do not thrive in urban environments, where one expects high levels of disturbance. One explanation for this pattern is that highly resilient species might not have been present in areas where humans settled. Indeed, humans have mostly settled around high primary productivity and relatively stable climate areas^39^. By offering stable environmental conditions, these locations might not have favoured adaptations to disturbances^40,41^. In fact, resilience to environmental disturbances is costly in environments where this resilience is not frequently needed^42^. Thus, maintaining some resistance aspect comes at the expense of other functions, leading to a reduced competitive ability in stable environments^43^. In this study, we focused on the impact of human activities and infrastructure via the Human Footprint^44^. While this includes some of the most important sources of human impact on biodiversity, such as land-use change and high population density, it does not include other pressures. Future research should consider other potential disturbance sources, such as harvesting, the presence of invasive species, pathogens, and climatic change.

The impacts of human pressure on the life-history traits of our examined populations are mostly negligible. Still, we find a higher ability to shrink in plants and longer generation times for long-mobility animals located closer to human activities. These results strengthen our hypothesis that species’ demographic properties mitigate the impact of human activities. Indeed, shrinkage increases the chances of survival either through a reduction in respiratory demands^45^ and/or because it allows the population to persist in disturbed and unsuitable environments while waiting for better conditions to reinitiate growth^46^. A shorter time to maturity can also allow the production of more phenotypes better adapted to changing environments^47^. Moreover, even though we found no clear evidence that human activities impact reproductive rates, such effects have been demonstrated in other studies^48,49^. This emphasises the need for joint macroecological research, such as ours, with long-term monitoring, and experiment-based studies, to better capture all the aspects of the ecology and evolution of life. Joint macroecological studies would also be better able to mitigate the biases that often exist in large-scale studies or databases. Effectively, in our case, despite the value of the data contained in COMPADRE and COMADRE databases, geographic and taxonomic biases remain^50^. For instance, our data are not fully taxonomically representative; 279 species of plants represent close to 0.080% of all plants and 45 species of animals <0.003% within that kingdom, moreover these species are not equally distributed across the tree of life (Table S4). Additionally, we did not incorporate species’ population density or climate change effects within our models, even though recent studies have shown important effects of the human footprint on mammal population density^51^ and of demographic responses in both plants and animals to climate change^52,53^.

Our work highlights the importance of studying short-term metrics of performance in ecological systems. The recently developed resilience framework, founded on short-term (*i.e*., transient) demographic metrics^21,54^, allows us to explore immediate responses of species populations to disturbances. Surprisingly, these metrics are not yet commonly used in population ecology^21,55^. Despite this oversight, they remain a powerful instrument to inform conservation programs and species risk assessments. We also demonstrate that resilience strategies are becoming less and less diverse as human pressure increases, which is an overlooked form of biodiversity loss i.e. the loss of “adaptive potential”. This loss presents a risk to the survival of species less able to cope with the diversity of disturbances predicted in the near future^56^.

## Materials and Methods

### Demographic data provenance and selection

To obtain high-resolution population-structured data on species demography, we used the COMPADRE Plant Matrix Database v.5.0.0^22^ and the COMADRE Animal Matrix Database v.3.0.0^23^. These databases contain 9,121 matrix population models (MPMs) of 760 plant species (Table S2), and 2,277 MPMs of 446 animal species, respectively (Table S1). MPMs are discrete time structured population models in which the vital rates (e.g., survival, reproduction) are explicitly examined by classifying individuals into discrete stages^57^. We then calculated transient dynamics metrics that relate to key demographic resilience quantities^21^, vital rates^57^, and life-history traits^58^. These calculations used the mean MPM under unmanipulated conditions for a given population to summarise the dynamics of each of our 921 populations across their examined period (See Supplementary Material for more details about the demographic metrics estimated).

To assure the validity of our demographic data to for our comparative analyses we impose a series of selection criteria on the total of 11,398 MPMs in versions used of COMPADRE and COMADRE. We first removed species whose MPMs contain at least one stage-specific vital rate value of NA, as we needed information across the full lifecycle. Next, we retained only terrestrial populations with full geographic representation in the HFP database^24^. We also discarded MPMs whose dominant eigenvalue population growth rate (*λ*) ≥ 3, as those implied unrealistic growth potentials. To further ensure fair comparisons across species in their natural conditions, we also removed MPMs from populations that were subject to manipulations (*e.g*., prescribed burning regimes). Finally, we only kept MPMs that are ergodic (whose asymptotic dynamics are independent of the initial population structure), primitive (MPMs consisting of non-negative elements), and irreducible (all life cycle stages are either directly, or indirectly connected to one another) ^59,60^ using the R package popdemo^61^. This set of criteria resulted in 796 MPM from 279 plant species, 109 MPMs from 45 animal species, 10 MPM of one algal species and 6 MPM of one fungal species.

To test H3, we derived a set of life-history traits from the selected MPMs. However, as life-history traits are known to be partly correlated^58,62,63^, we checked for colinearity using Pearson correlation tests (with a threshold of 0.6) with the cor() function in R version 4.0.1^64^.

### Resilience framework

To understand how human presence shapes the resilience strategies of species, we examined the volume occupied by our populations in a 4D space describing resistance, speed of recovery, compensation, and human pressure. Resistance, speed of recovery, and compensation were calculated as the inverse of the first step attenuation (*<u>ρ</u>_1_ = minCS*(Â)), the log of the damping ratio (ρ = *λ*_1_/∥*λ*_2_∥) and the log of the maximum amplification 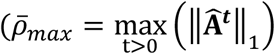. Where ***Â*** is the normalised ***A*** population matrix, which involves scaling each element of ***A*** by *λ*_1_, *minCS* is the minimum column sum of a matrix, *λ*_1_ is the dominant eigenvalue, *λ*_2_ is the subdominant eigenvalue. To test the distribution of these metrics we used the method described in Diaz et al.^65^. Briefly, this approach consists of quantifying convex hulls based on the empirical distribution of the resistance, recovery time, and compensation of our populations, compared to different null model where these traits are (1) uniformly distributed between the maximum and the minimum values of the observations, (2) normally distributed with the mean and the variance of the normal distribution being equal to the mean and the variance of the observations respectively, and (3) gamma distributed along the axis of human presence. To model the gamma distribution, we used the following parameters:

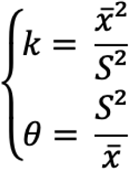

with *k* being the shape and *²* the scale of the gamma distribution, *<u>x</u>* the mean of the observed data and *S^2^* the variance of the observed data.

### Human pressure data

To quantify human pressure, we used the Human Footprint (HFP)^24^, which has been shown to correlate strongly with species extinction risk^66,67^. The HFP is composed of eight layers that cover all terrestrial surfaces of the globe, excluding Antarctic land areas, at a 1km^²^ resolution. The HFP allows us to determine at a high spatial resolution not only the overall effects of humans on the demographic resilience and life-history traits of natural populations, but also their underlying putative causes (e.g., density of built infrastructure, proximity to railways). The HFP assigns a score of 50 based on multiple human activities on the environment at each pixel (where a pixel is 1km^²^ x 1km^²^). These eight layers of activity include human population density, roads, navigable waterways, railways, night-time lights, built environment—which is the density of buildings and paved land estimated through satellite images, pasture lands, and croplands (Fig. 1). Two sets of maps are available in the HFP, for 1993 and 2009. These two years allow for a more comprehensive fit between the demographic data used here (below) and the human pressure at the time of the record. Specifically, for each demographic record, we selected the HFP record that was closest to the year when the demographic data were collected.

We used multivariate analyses to assign a human-pressure score to each of the natural populations we assessed—this approach allowed us to account for heterogeneity in the human activities and their different potential effects. We first performed a principal component analysis (PCA) on all eight HFP layers. To do so, we rescaled the HFP map at a 10 km^2^ × 10 km^2^ resolution, sub-sampling just one pixel in each grid cell. Next, we extracted 60,000 random data points from the 10 km^²^ map and used that subset of data to run the PCA analysis. Because the HFP data were linked to the 1993 or the 2009 HFP version, we compared the PCA outputs coming only from the 1993 map, from the 2009 map, or from a mix of both. Finally, we finally decided to use the subset of 60,000 random but unique points distributed around the map and the two HFP dates.

### Phylogenetic information gathering

To quantify and explicitly consider the effect of potential phylogenetic co-variation in traits in our analyses, we constructed two phylogenetic trees resolved down to the population level, one for plants (see Supplementary Material), and another one for animals (see Supplementary Material). We used Timetree^68^ to create an animal phylogeny to accommodate our 109 populations from 45 examined animal species, and the package V.phylomaker^69^ to create a plant phylogeny to accommodate our 796 populations from 279 examined plant species. As some sources in COMADRE and COMPADRE contain multiple populations per species, we created our phylogenies at the population level instead of the species level. To do so, each population from the same species was separated by a small distance (*ε*= 0.0000001 normalized units), thus assuming that populations within the same species are closely related. As the order of the population divergences within our phylogenetic analyses might have an impact on the measure of the phylogenetic signal and on the phylogenetic correction within our future analyses, we ran a sensitivity analysis of some phylogenetic signals using different trees with a random order of the populations within the tree. These sensitivity analyses showed no effect (See Supplementary Material).

### Demographic traits regression

We developed two models of regression to study the interaction between human activities and species’ demographic resilience components. For both models, human activities were described according to the two dominant axes of variation in human activities estimated through the PCA. In both models, we controlled for trait variation due to the size of individuals (*i.e*., adult body mass for animals, maximum height for plants), spatial autocorrelation, the phylogenetic or evolutionary history autocorrelation, and the complexity of the life-history that was estimated via the dimensions of the relevant MPM. Importantly here, it has been shown that life-history traits scale with size in animals, especially in mammals^70,71^. Similarly, it has been shown that spatially close populations are more likely to be similar to distant populations than expected by chance^72,73^.

Our first model to test the effects of human pressure on demographic traits was based on the study of the residuals after removing the confounding effects: animal size/plant height, spatial autocorrelations, phylogenetic relationships. We did this to account for autocorrelations due to location and ancestry^72,74^. We incorporated a correction for body mass and plant height (later called size) in our study when linearity was assessed in the relationship between the trait and the size. Next, we assessed the effect of spatial autocorrelation on our models by calculating the Moran’s index associated with the different traits or with the residuals of the regression of the trait and size when a size effect was detected. If spatial autocorrelation was found, we incorporated a correction for the location in the model^75^. To do so, we used the tensor function of generalized additive models (gam) with longitude and latitude^76,77^ using R-package mgcv version 1.8-28 (*Equation 1*). Finally, we ran a Markov Chain Monte Carlo generalized linear mixed model (MCMCglmm) of the residuals of the traits corrected (or not) for size and spatial autocorrelation on the HFP PCA main dimensions and that considered the phylogeny of the populations (*Equation 2*). Phylogenetic corrections for plant and animal populations were undertaken depending on the value of Pagels’ *λ*^78^ (Pagels’ *λ* significantly different of 0).

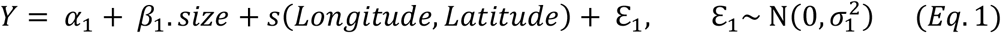

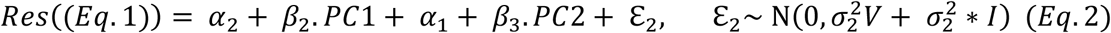

Where Y is the trait of interest, PC1 and PC2 are the two first axis of the HFP PCA, Res(Eq.1) are the residuals of the gam models in equation 1, V is the covariance matrix derived from the phylogeny and 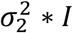 gives the non-phylogenetic residual variance.

Our second series of models to validate the new model developed aimed to disentangle spatial and phylogenetic corrections. As working on residuals can be controversial because one might neglect important pieces of information^79,80^—namely that model estimates based on residuals can provide biased estimates and this approach also increases the significance of variables that are more sensitive to this method—we ran models with the same original data but without using residuals. We included size as an explanatory factor when it was relevant (Fisher’s exact test) and ran a Markov Chain Monte Carlo generalized linear mixed model (MCMCglmm) to account for evolutionary history and a Generalized Additive Models (gam) model with longitude and latitude to account for spatial distribution.

All the analyses were performed in R version 4.0.1^64^.

## Acknowledgement

We are grateful to the SalGo team for their valuable comments, help, and discussions. We also acknowledge the use of and support from the University of Oxford’s Advanced Research Computing (ARC)._PC was supported by a Ramón Areces Foundation postdoctoral scholarship (BEVP30P01A5816) hosted by RSG. MDM was supported by the MUR Rita Levi Montalcini program. RS-G was supported by a NERC Independent Research Fellowship (NE/M018458/1).

