## Supplementary Online Materials for "Human disturbances erode the diversity of species resilience strategies"

### Appendix S1: Extended methods

#### Demographic data selection

To run our analyses regarding the human-activity correlates of natural population resilience, we carefully selected a subset of species from peer-reviewed publications archived in the COMPADRE Plant Matrix Database[1] (version 5.0.0) and the COMADRE Animal Matrix database[2] (version 3.0.0) . The species and original sources are shown in Table S1 and S2. To allow for inter-specific comparisons, we imposed a series of criteria on the matrix population models (MPMs) stored in COMPADRE and COMADRE. First, only terrestrial species were considered, as the Human Footprint index is only available for terrestrial habitats[3]. Second, we retained only studies for which we were able to obtain a mean matrix population model that would describe the dynamics of the species over its whole duration, thus removing exceptional seasonal dynamics. Third, we only considered MPMs in unmanipulated conditions (*i.e.* no experimental manipulations) to link transient metrics to human pressures and avoid artefacts resulting from human manipulation. Fourth, we discarded MPMs with abnormally high values of population growth rate ($\lambda > 3$) as they indicate unrealistic or exceptional conditions. Finally, we removed MPMs that were not ergodic, primitive and irreducible, as these conditions are required to derive most of our life history traits and transient dynamics' metrics[4].

**Table S1.** List of the animal species used in this study, from the COMADRE Animal Matrix Database. # pop indicates the number of populations we had for each population.

| Latin name | Family | Common name | # pop | Authors | DOI.ISBN | Publication year |
|---|---|---|---|---|---|---|
| *Acinonyx jubatus* | Felidae | Cheetah | 1 | Lubben; Tenhumberg; Tyre; Rebarber | 10.1016/j.biocon.2007.11.003 | 2008 |
| *Alces alces* | Cervidae | Moose | 4 | Carroll, Kvalnes; Saether; Haanes; Røed; Engen; Solberg | NA, 10.1111/evo.12952 | 2013, 2016 |
| *Ammodramus savannarum* | Emberizidae | Grasshopper sparrow | 2 | Hovick; Miller | 10.1007/s10980-013-9896-7 | 2013 |
| *Anser anser* | Anatidae | Greylag goose | 1 | Klok; van Turnhout; Willems; Voslamber; Ebbinge; Schekkerman | 10.1163/157075610X523260 | 2010 |
| *Anthropoides paradiseus* | Gruidae | Blue crane | 1 | Altwegg; Anderson | 10.1111/j.1365-2435.2009.01563.x | 2009 |
| *Bonasa umbellus* | Phasianidae | Ruffed grouse | 7 | Tirpak; Giuliano; Miller; Allen; Bittner; Buehler; Edwards; Harper; Igo; Norman; Seamster; Stauffer | 10.1016/j.biocon.2006.06.014 | 2006 |
| *Bostrychia hagedash* | Threskiornithidae | Hadeda ibis | 1 | Duckworth; Altwegg; Harebottle | 10.1007/s10336-011-0758-2 | 2011 |
| *Calidris temminckii* | Scolopacidae | Baltic temminck's stint | 2 | Koivula; Pakanen; Rönkä; Belda | 10.1111/j.0908-8857.2008.04189.x | 2008 |
| *Callospermophilus lateralis* | Sciuridae | Golden-mantled ground squirrel | 1 | Hostetler; Kneip; Van Vuren; Oli | 10.1371/jourNAl.pone.0034379 | 2012 |
| *Calyptorhynchus lathami* | Psittacidae | Glossy Black Cockatoo | 2 | Harris; Fordham; Mooney; Pedler; Araújo; Paton; Stead; Watts; Akçakaya; Brook | 10.1111/j.1365-2664.2012.02163.x | 2012 |

| | | | | | | |
|---|---|---|---|---|---|---|
| *Cebus capucinus* | Cebidae | White faced capuchin monkey | 1 | Morris; Altmann; Brockman; Cords; Fedigan; Pusey; Stoinski; Bronikowski; Alberts; Strier | 10.1086/657443 | 2011 |
| *Cervus elaphus* | Cervidae | Elk | 2 | Hebblewhite; Merrill | 10.1111/j.1600-0706.2011.19436.x | 2011 |
| *Chelodina expansa* | Chelidae | Broad-shelled turtle | 2 | Spencer; Thomson | 10.1111/j.1523-1739.2005.00487.x | 2005 |
| *Cryptophis nigrescens* | Elapidae | Common small-eyed snake | 1 | Webb; Brook; Shine | 10.1046/j.1440-1703.2002.00463.x | 2002 |
| *Dasypus novemcinctus* | Dasypodidae | Nince-banded armadillo | 3 | Oli; Loughry; Caswell; Perez-Heydrich; McDonough; Truman | 10.1016/j.ecolmodel.2017.02.001 | 2017 |
| *Eidolon helvum* | Pteropodidae | Straw-colored fruit bat | 1 | Hayman; McCrea; Restif; Suu-Ire; Fooks; Wood; Cunningham; Rowcliffe | 10.1017/S0950268812000167 | 2012 |
| *Emydura macquarii* | Chelidae | Macquarie turtle | 2 | Spencer; Thomson | 10.1111/j.1523-1739.2005.00487.x | 2005 |
| *Erythrocebus patas* | Cercopithecidae | Patas monkey | 1 | Isbell; Young; Jaffe; Carlson; Chancellor | 10.1007/s10764-009-9332-7 | 2009 |
| *Falco naumanni* | Falconidae | Lesser kestrel | 1 | Hiraldo; Negro; Donazar; Gaona | 10.2307/2404688 | 1996 |
| *Falco peregrinus* | Falconidae | Peregrine falcon | 1 | Altwegg; Jenkins; Abadi | 10.1111/ibi.12125 | 2013 |
| *Forpus passerinus* | Psittacidae | Green-rumped parrotlets | 2 | Sandercock; Beissinger | 10.1080/02664760120108818 | 2002 |
| *Fulmarus glacialis* | Procellariidae | Northern fulmar | 1 | Kerbiriou; Le Viol; Bonnet; Robert | 10.1007/s10144-012-0306-9 | 2012 |
| *Gavia immer* | Gaviidae | Great northern diver | 2 | Grear; Meyer; Cooley; Kuhn; Piper; Mitro; Vogel; Taylor; Kenow; Craig; Nacci | 10.2193/2008-093 | 2010 |

| | | | | | | |
|---|---|---|---|---|---|---|
| *Giraffa camelopardalis* | Giraffidae | Giraffe | 2 | Strauss; Kilewo; Rentsch; Packer | 10.1007/s10144-015-0499-9 | 2015 |
| *Himantopus novaezelandiae* | Recurvirostridae | Black stilt | 1 | Cruz; Pech; Seddon; Cleland; Nelson; Sanders; Maloney | 10.1016/j.biocon.2013.09.005 | 2013 |
| *Hoplocephalus bungaroides* | Elapidae | Broad-headed snake | 1 | Webb; Brook; Shine | 10.1046/j.1440-1703.2002.00463.x | 2002 |
| *Lagopus leucura* | Phasianidae | White-tailed ptarmigan | 2 | Wilson; Martin | 10.1186/1472-6785-12-9 | 2012 |
| *Lagopus muta* | Phasianidae | Rock ptarmigan | 2 | Wilson; Martin | 10.1186/1472-6785-12-9 | 2012 |
| *Malaclemys terrapin* | Emydidae | Diamondback terrapin | 1 | Crawford; Maerz; Nibbelink; Buhlmann; Norton | 10.1111/1365-2664.12194 | 2013 |
| *Odocoileus virginianus* | Cervidae | White-tailed deer | 13 | Chitwood; Lashley; Kilgo; Moorman; Deperno, Edmunds | 10.1002/jwmg.835, NA | 2015, 2013 |
| *Onychogalea fraenata* | Macropodidae | Bridled nailtail wallaby | 1 | Fisher; Hoyle; Blomberg | 10.2307/2641054 | 2000 |
| *Ovis canadensis* | Bovidae | Bighorn sheep | 9 | Rubin; Boyce; Caswell-Chen, Coulson; Gaillard; Festa-Bianchet | 10.2307/3803144, 10.1111/j.1365-2656.2005.00975.x | 2002, 2005 |
| *Panthera pardus* | Felidae | Leopard | 1 | Balme; Slotow; Hunter | 10.1016/j.biocon.2009.06.020 | 2009 |
| *Phrynosoma cornutum* | Phrynosomatidae | Texas horned lizard | 2 | Wolf; Hellgren; Schauber; Bogosian III; Kazmaier; Ruthven III; Moody | 10.1007/s10144-014-0450-5 | 2014 |
| *Podocnemis expansa* | Podocnemididae | Arrau turtle | 1 | Mogollones; Rodríguez; Hernández; Barreto | 10.2744/CCB-0778.1 | 2010 |
| *Propithecus edwardsi* | Indriidae | Milne-Edwards' sifaka | 2 | Dunham; Erhart; Overdorff; Wright | 10.1016/j.biocon.2007.10.006 | 2008 |

| | | | | | | |
|---|---|---|---|---|---|---|
| *Sceloporus arenicolus* | Phrynoso matidae | Dunes sagebrush lizard | 7 | Ryberg; Hill; Painter; Fitzgerald | 10.1111/cobi.12429 | 2014 |
| *Sceloporus grammicus* | Phrynoso matidae | Mesquite lizard | 9 | Pérez-Mendoza, Zuniga-Vega; Mendez-dela Cruz; Cuellar | 10.1655/HERPETOLOGICA-D-12-00038R2, 10.1139/Z08-124 | 2013, 2008 |
| *Setophaga cerulea* | Parulidae | Cerulean warbler | 1 | Jones; Barg; Sillett; Veit; Robertson | 10.1642/0004-8038(2004)121[0015:MEOSAP]2.0.CO;2 | 2004 |
| *Sterna hirundo* | Laridae | Common tern | 2 | Szostek | 10.1007/s10336-011-0745-7 | 2011 |
| *Tamiasciurus hudsonicus* | Sciuridae | American red squirrel | 1 | McAdam; Boutin; Sykes; Humphries | 10.2980/1195-6860(2007)14[362:LHOFRS]2.0.CO;2 | 2007 |
| *Turdus torquatus* | Turdidae | Ring ouzel | 3 | Sim; Rebecca; Ludwig; Grant; Reid | 10.1111/j.1365-2656.2010.01750.x | 2011 |
| *Urocyon littoralis* | Canidae | Island fox | 2 | Hudgens; Garcelon | 10.1007/s00442-010-1761-7 | 2011 |
| *Ursus americanus* | Ursidae | American black bear | 3 | Lewis; Breck; Wilson; Webb, Hebblewhite; Percy; Serrouya | 10.1016/j.ecolmodel.2014.08.021, 10.1016/S0006-3207(02)00341-5 | 2014, 2003 |
| *Vipera aspis* | Viperidae | Asp viper | 1 | Altwegg; Dummermuth; Anholt; Flatt | 10.1111/j.0030-1299.2001.13723.x | 2005 |

**Table S2.** List of the plant species used in this study, from the COMPADRE Plant Matrix Database. # pop indicates the number of populations we had for each population.

| Latin name | Family | Common name | # pop | Authors | DOI or ISBN | Publication year |
|---|---|---|---|---|---|---|
| *Acacia bilimekii* | Leguminosae | NA | 1 | Jiménez-Lobato; Valverde | 10.1016/j.jaridenv.2005.07.002 | 2006 |
| *Acacia suaveolens* | Leguminosae | NA | 1 | Warton; Wardle | 10.1046/j.1442-9993.2003.01246.x | 2003 |
| *Acer palmatum* | Sapindaceae | NA | 1 | Tanaka; Shibata; Masaki; Iida; Niiyama; Abe; Kominomi; Nokashizuka | 10.3170/2007-8-18342 | 2008 |
| *Acer pictum* | Sapindaceae | NA | 1 | Tanaka; Shibata; Masaki; Iida; Niiyama; Abe; Kominomi; Nokashizuka | 10.3170/2007-8-18342 | 2008 |
| *Acer rufinerve* | Sapindaceae | NA | 1 | Tanaka; Shibata; Masaki; Iida; Niiyama; Abe; Kominami; Nokashizuka | 10.3170/2007-8-18342 | 2008 |
| *Actaea elata* | Ranunculaceae | NA | 1 | Mayberry; Elle | 10.1007/s00442-010-1809-8 | 2010 |
| *Adenocarpus aureus gibbsianus* | Leguminosae | NA | 1 | Iriondo; Albert; Giménez; Lozano; Escudero | 978-84-8014-746-0 | 2009 |
| *Adesmia volckmannii* | Leguminosae | NA | 1 | Cipriotti; Aguiar | 10.1111/j.1654-109X.2011.01138.x | 2012 |
| *Aechmea magdalenae* | Bromeliaceae | NA | 3 | Ticktin; Nantel | 10.1016/j.biocon.2004.03.019 | 2004 |
| *Agrimonia eupatoria* | Rosaceae | NA | 3 | Kiviniemi | 10.1111/j.1523-1739.2011.01691.x | 2002 |

| | | | | | | |
|---|---|---|---|---|---|---|
| *Alliaria petiolata* | Brassicaceae | Garlic mustard | 10 | Evans; Davis; Raghu; Ragavendran; Landis; Schemske | 10.1890/11-1291.1 | 2012 |
| *Anarrhinum fruticosum* | Plantaginaceae | NA | 1 | Iriondo; Albert; Giménez; Lozano; Escudero | 978-84-8014-746-0 | 2009 |
| *Androsace elongata* | Primulaceae | NA | 4 | Dostál | 10.1111/j.1654-1103.2007.tb02519.x | 2007 |
| *Anemone patens* | Ranunculaceae | Prairie crocus | 1 | Williams; Crone | 10.1890/0012-9658(2006)87[3200:TIOIGO]2.0.CO;2 | 2006 |
| *Anthericum ramosum* | Asparagaceae | NA | 1 | Černá; Münzbergová | 10.1371/journal.pone.0075563 | 2013 |
| *Anthyllis vulneraria L.* | Fabaceae | NA | 20 | Davison; Jacquemyn; Adriaens; Honnay; de Kroon; Tuljapurkar | NA | 2010 |
| *Antirrhinum molle lopesianum* | Acanthaceae | NA | 1 | Iriondo; Albert; Giménez; Lozano; Escudero | 978-84-8014-746-0 | 2009 |
| *Antirrhinum subbaeticum* | Plantaginaceae | NA | 2 | Iriondo; Albert; Giménez; Lozano; Escudero | 978-84-8014-746-0 | 2009 |
| *Aquilaria crassna* | Thymelaeaceae | NA | 1 | Zhang; Brockelman; Allen | 10.1016/j.biocon.2008.04.015 | 2008 |
| *Aquilegia chrysantha* | Ranunculaceae | NA | 1 | Stubben | NA | 2007 |
| *Araucaria muelleri* | Araucariaceae | NA | 4 | Enright; Miller; Perry; Goldblum; Jaffré | 10.1111/aec.12045 | 2014 |
| *Arenaria grandiflora bolosii* | Caryophyllaceae | NA | 1 | Iriondo; Albert; Giménez; Lozano; Escudero | 978-84-8014-746-0 | 2009 |
| *Arenaria serpyllifolia* | Caryophyllaceae | NA | 4 | Dostál | 10.1658/1100-9233(2007)18[91:PDOAIP]2.0.CO;2 | 2007 |

| | | | | | | |
|---|---|---|---|---|---|---|
| *Argyroxiphium sandwicense* | Composita e | Haleakala silversword | 1 | Forsyth | 10.1007/s00442-003-1295-3 | 2003 |
| *Armeria merinoi* | Plumbagin aceae | NA | 2 | Iriondo; Albert; Giménez; Lozano; Escudero | 978-84-8014-746-0 | 2009 |
| *Arnica angustifolia* | Composita e | NA | 1 | Jakalaniemi | 10.1016/j.envexpbot.2011.03.0 13 | 2011 |
| *Artemisia genipi* | Composita e | NA | 2 | Marcante; Winkler; Erschbamer | 10.1093/aob/mcp047 | 2009 |
| *Asarum canadense* | Aristolochi aceae | Wild ginger | 7 | Damman; Cain | 10.1046/j.1365-2745.1998.00242.x | 1998 |
| *Asclepias meadii* | Apocynac eae | Mead's milkweed | 2 | Bell; Bowles; McEachern | 978-3-642-07869-9 | 2003 |
| *Aspasia principissa* | Orchidace ae | NA | 1 | Zotz; Schmidt | 10.1016/j.biocon.2005.07.022 | 2006 |
| *Asplenium adulterinum* | Aspleniac eae | NA | 7 | Bucharová; Münzbergová; Tájek | 10.3732/ajb.0900351 | 2010 |
| *Asplenium cuneifolium* | Aspleniac eae | NA | 5 | Bucharova; Munzbergova; Tajek | 10.3732/ajb.0900351 | 2010 |
| *Asplenium scolopendrium* | Aspleniac eae | NA | 1 | Bremer; Jongejans | 10.1007/s10144-009-0143-7 | 2010 |
| *Astragalus alopecurus* | Legumino sae | NA | 5 | Nicolè | NA | 2005 |
| *Astragalus cottonii* | Fabaceae | NA | 3 | Kaye | NA | 1990 |
| *Astragalus scaphoides* | Legumino sae | NA | 2 | Lesica | NA | 1995 |
| *Astragalus tremolsianus* | Legumino sae | NA | 1 | Iriondo; Albert; Giménez; Lozano; Escudero | 978-84-8014-746-0 | 2009 |
| *Astragalus tyghensis* | Legumino sae | NA | 5 | Kaye; Pyke | 10.1890/0012-9658(2003)084[1464:TEOSTO ]2.0.CO;2 | 2003 |
| *Astrophytum asterias* | Cactaceae | NA | 4 | Martinez-Avalos | NA | 2007 |

| | | | | | | |
|---|---|---|---|---|---|---|
| *Astrophytum capricorne* | Cactaceae | NA | 1 | Mandujano; Bravo; Verhulst; Carrillo-Angeles; Golubov | 10.1016/j.actao.2014.12.004 | 2015 |
| *Atriplex acanthocarpa* | Amaranthaceae | Tubercled salt-bush | 1 | Verhulst; Montaña; Mandujano; Franco | 10.1007/s00442-008-0980-7 | 2008 |
| *Atriplex canescens* | Amaranthaceae | Four-winged salt-bush | 1 | Verhulst; Montaña; Mandujano; Franco | 10.1007/s00442-008-0980-7 | 2008 |
| *Balsamorhiza sagittata* | Compositae | Arrowleaf balsamroot | 1 | Crone; Marler; Pearson | 10.1111/j.1365-2664.2009.01635.x | 2009 |
| *Bertholletia excelsa* | Lecythidaceae | Brazil nut tree | 4 | Zuidema; Boot | 10.1017/S0266467402002018 | 2002 |
| *Boechera fecunda* | Brassicaceae | NA | 3 | Lesica; Shelly | 10.2307/2445615 | 1995 |
| *Boltonia decurrens* | Compositae | NA | 1 | Smith; Caswell; Mettler-Cherry | 10.1890/04-0434 | 2005 |
| *Borassus aethiopum* | Arecaceae | NA | 3 | Barot; Gignoux; Vuattoux | 10.1017/S0266467400001620 | 2000 |
| *Brassica insularis* | Brassicaceae | NA | 4 | Noel; Maurice; Mignot; Glémin; Carbonell; Justy; Guyot; Olivieri; Petit | 10.1007/s10592-010-0056-1 | 2010 |
| *Calamus nambariensis* | Arecaceae | NA | 1 | Binh | NA | 2009 |
| *Calamus rhabdocladus* | Arecaceae | NA | 1 | Binh | NA | 2009 |
| *Calathea micans* | Marantaceae | NA | 2 | Le Corff; Horvitz | 10.1016/j.ecolmodel.2005.05.009 | 2005 |
| *Calocedrus decurrens* | Cupressaceae | NA | 1 | van Mantgem; Stephenson | 10.1111/j.1365-2745.2005.01007.x | 2005 |
| *Calochortus lyallii* | Liliaceae | NA | 20 | Miller; Antos; Allen | NA | 2004 |
| *Calochortus macrocarpus* | Liliaceae | NA | 5 | Miller; Antos; Allen | NA | 2004 |

| | | | | | | |
|---|---|---|---|---|---|---|
| *Carduus nutans* | Compositae | Nodding thistle; Musk thistle | 4 | Jongejans; Sheppard; Shea | 10.1111/j.1365-2664.2006.01228.x | 2006 |
| *Carex humilis* | Cyperaceae | NA | 3 | Wikberg; Svensson | 10.1007/s11258-005-9006-2 | 2006 |
| *Carlina vulgaris* | Asteraceae | NA | 4 | Jongejans; Jorritsma-Wienk; Becker; Dostál; Mildén | 10.1111/j.1365-2745.2009.01612.x | 2010 |
| *Castanea dentata* | Fagaceae | American chestnut | 3 | Davelos; Jarosz | 10.1111/j.0022-0477.2004.00907.x | 2004 |
| *Catopsis compacta* | Bromeliaceae | NA | 1 | del Castillo; Trujillo-Argueta; Rivera-Garcia; Gómez-Ocampo; Mondragón-Chaparro | 10.1002/ece3.765 | 2013 |
| *Cecropia obtusifolia* | Urticaceae | NA | 1 | Alvarez-Buylla | 10.1086/285599 | 1994 |
| *Centaurea corymbosa* | Asteraceae | NA | 6 | Belaid; Maurice; Fréville; Carbonell; Imbert | 10.1016/j.biocon.2018.04.019 | 2018 |
| *Centaurea horrida* | Compositae | NA | 2 | Pisanu; Farris; Filigheddu; Begona Garcia | 10.1007/s11258-012-0110-9 | 2012 |
| *Cerastium pumilum* | Caryophyllaceae | European chickweed | 1 | Burns; Pardini; Schutzenhofer; Chung; Seidler; Knight | 10.1890/12-1310.1 | 2013 |
| *Chamaecrista lineata keyensis* | Leguminosae | NA | 6 | Liu; Menges; Quintana-Ascencio | 10.1890/03-5382 | 2005 |
| *Chamaedorea elegans* | Arecaceae | NA | 1 | Valverde; Hernandez-Apolinor; Mendoza-Amarom | 10.1300/J091v23n01_05 | 2006 |
| *Chamaedorea radicalis* | Arecaceae | NA | 5 | Endress; Gorchov; Robert; Noble, Berry; Gorchov; Endress; Stevens | 10.1890/02-5365, 10.1007/s10144-007-0067-z | 2004, 2008 |

| | | | | | | |
|---|---|---|---|---|---|---|
| *Cheirolophus metlesicsii* | Composita e | NA | 1 | Iriondo; Albert; Giménez; Lozano; Escudero | 978-84-8014-746-0 | 2009 |
| *Cirsium dissectum* | Composita e | Meadow thistle | 4 | Jongejans; de Vere; de Kroon | 10.1007/s11258-008-9397-y | 2008 |
| *Cirsium perplexans* | Composita e | Rocky Mountain thistle | 2 | Dodge | NA | 2005 |
| *Cirsium pitcheri* | Composita e | Pitcher's thistle | 7 | Bell; Bowles; McEachern, Ellis; Williams; Lesica; Bell; Bierzychudek; Bowles; Crone; Doak; Ehrlén; Ellis-Adam; McEachern; Ganesan; Latham; Luijten; Kaye; Knight; Menges; Morris; Den Nijs; Oostermeijer; Quintana-Ascencio; Shelly; Stanley; Thorpe; Ticktin; Valverde; Weekley, Bell; Powell; Bowles, Jolls; Marik; Hamze; Havens | 978-3-642-07869-9, 10.1890/11-1052.1, 10.1002/jwmg.525, 10.1016/j.biocon.2015.04.006 | 2003, 2012, 2013, 2015 |
| *Cirsium vulgare* | Composita e | Bull thistle; Spear thistle | 3 | Bullock; Hill; Silvertown | 10.2307/2261390 | 1994 |
| *Clidemia hirta* | Melastom ataceae | NA | 2 | DeWalt | 10.1007/s10530-005-5277-8 | 2006 |
| *Cornus florida* | Cornacea e | Eastern flowering dogwood | 1 | Vejdani | NA | 2006 |
| *Cryptantha flava* | Boraginac eae | NA | 2 | Lucas; Forseth; Casper, Salguero-Gomez; Kempenich; Forseth; Casper | 10.1111/j.1365-2745.2007.01350.x, 10.1890/13-1256.1 | 2008, 2014 |

| | | | | | | |
|---|---|---|---|---|---|---|
| *Cypripedium calceolus* | Orchidaceae | Lady slipper | 6 | García; Goñi; Guzman, Nicolè; Brzosko; Till-Bottraud | 10.1111/j.1523-1739.2010.01466.x, 10.1111/j.1365-2745.2005.01010.x | 2010, 2005 |
| *Cypripedium fasciculatum* | Orchidaceae | Clustered lady's slipper | 3 | Thorpe; Stanley; Kayne; Latham | NA | 2011 |
| *Cyrtandra dentata* | Gesneriaceae | Mountain cyrtandra | 18 | Bialic-Murphy; Gaoue; Kawelo | 10.1111/1365-2664.12868 | 2017 |
| *Cytisus scoparius* | Leguminosae | Common broom; Scotch broom; English broom | 1 | Neubert; Parker | 10.1111/j.0272-4332.2004.00481.x | 2004 |
| *Dactylorhiza lapponica* | Orchidaceae | NA | 2 | Sletvold; Øien; Moen | 10.1016/j.biocon.2009.12.017 | 2010 |
| *Daemonorops poilanei* | Arecaceae | NA | 1 | Binh | NA | 2009 |
| *Daphne rodriguezii* | Thymelaeaceae | NA | 3 | Rodriguez-Perez; Traveset | 10.1111/j.1600-0706.2011.19946.x | 2012 |
| *Dicentra canadensis* | Papaveraceae | Squirrel corn | 3 | Lin; Miriti; Goodell | 10.1002/ece3.2163 | 2016 |
| *Dicerandra frutescens* | Lamiaceae | Florida scrub mint | 7 | Menges; Quintana-Ascencio; Weekley; Gaoue | 10.1016/j.biocon.2005.08.002 | 2006 |
| *Dioon caputoi* | Zamiaceae | NA | 1 | Cabrera-Toledo | NA | 2009 |
| *Dioon merolae* | Zamiaceae | NA | 2 | Lázaro-Zermeño; González-Espinosa; Mendoza; Martinez-Ramos; Quintana-Ascencio | 10.1016/j.foreco.2010.10.028 | 2011 |
| *Dioon sonorense* | Zamiaceae | NA | 1 | Álvarez-Yépiz; Dovčiak; Búrquez | 10.1016/j.biocon.2010.08.007 | 2011 |
| *Dioscorea chouardii* | Dioscoreaceae | NA | 1 | Garcia | 10.1016/S0006-3207(01)00113-6 | 2003 |

| | | | | | | |
|---|---|---|---|---|---|---|
| *Dracocephalum austriacum* | Lamiaceae | Austrian dragonbead | 12 | Dostálek; Münzbergova | 10.1007/s12224-012-9132-2, NDY | 2013 |
| *Dypsis decaryi* | Arecaceae | Triangle plam | 3 | Ratsirarson; Silander; Richard | 10.1046/j.1523-1739.1996.10010040.x | 1996 |
| *Echinacea angustifolia* | Asteraceae | Narrow-leaved purple coneflower | 6 | Dykstra, Hurlburt | NA | 2013, 1999 |
| *Encephalartos cycadifolius* | Zamiaceae | NA | 1 | Raimondo; Donaldson | 10.1016/S0006-3207(02)00303-8 | 2003 |
| *Encephalartos villosus* | Zamiaceae | NA | 1 | Raimondo; Donaldson | 10.1016/S0006-3207(02)00303-8 | 2003 |
| *Entandrophragma cylindricum* | Meliaceae | Sapelli | 1 | Picard; Yalibanda; Namkosserena; Baya | 10.1016/j.foreco.2008.02.041 | 2008 |
| *Epilobium latifolium* | Onagraceae | Dwarf fireweed | 1 | Doak | 10.2307/1941457 | 1992 |
| *Epipactis atrorubens* | Orchidaceae | Darkred helleborine | 1 | Hens; Pakanen; Jäkäläniemi; Tuomi; Kvist | 10.1016/j.biocon.2017.04.019 | 2017 |
| *Eremospatha macrocarpa* | Arecaceae | NA | 2 | Kouassi; Barot; Gignoux; Bi | 10.1017/S0266467408005312 | 2008 |
| *Eriogonum longifolium gnaphalifolium* | Polygonaceae | Buckwheat | 13 | Satterthwaite; Menges; Quintana-Ascencio | 10.1890/1051-0761(2002)012[1672:ASBPVI]2.0.CO;2 | 2002 |
| *Eritrichium caucasicum* | Boraginaceae | NA | 2 | Logofet; Kazantseva; Belova; Onipchenko | 10.1134/S2079086418030076 | 2018 |
| *Erodium paularense* | Geraniaceae | NA | 1 | Iriondo; Albert; Giménez; Lozano; Escudero | 978-84-8014-746-0 | 2009 |
| *Eryngium alpinum* | Apiaceae | NA | 1 | Andrello; Bizoux; Barbet-Massin; Gaudeul; Nicolè; Till-Bottraud | 10.1016/j.biocon.2011.12.012 | 2012 |
| *Eryngium cuneifolium* | Apiaceae | Florida rosemary scrub | 8 | Menges; Quintana-Ascencio | 10.1890/03-4029 | 2004 |

| | | | | | | |
|---|---|---|---|---|---|---|
| *Eryngium maritimum* | Apiaceae | NA | 1 | Curie; Stabbetorp; Nordal | 10.1111/j.1756-1051.2004.tb01647.x | 2007 |
| *Escobaria robbinsorum* | Cactaceae | NA | 4 | Schmalzel; Reichenbacher; Rutman | NA | 1995 |
| *Escontria chiotilla* | Cactaceae | NA | 2 | Ortega-Baes | NA | 2001 |
| *Eupatorium perfoliatum* | Compositae | Common boneset | 3 | Byers; Meagher | 10.1890/1051-0761(1997)007[0519:ACODCI]2.0.CO;2 | 1997 |
| *Eupatorium resinosum* | Compositae | Pine Barrens boneset | 1 | Byers; Meagher | 10.1890/1051-0761(1997)007[0519:ACODCI]2.0.CO;2 | 1997 |
| *Euphorbia fontqueriana* | Euphorbiaceae | NA | 1 | Iriondo; Albert; Giménez; Lozano; Escudero | 978-84-8014-746-0 | 2009 |
| *Euterpe edulis* | Arecaceae | NA | 1 | Silva-Matos; Freckleton; Watkinson | 10.1890/0012-9658(1999)080[2635:TRODDI]2.0.CO;2 | 1999 |
| *Euterpe oleracea* | Arecaceae | NA | 2 | Arango; Duque; Muñoz | 10.15517/rbt.v58i1.5222 | 2010 |
| *Euterpe precatoria* | Arecaceae | NA | 2 | Otárola; Avalos | 10.3732/ajb.1400089 | 2014 |
| *Festuca eskia* | Poaceae | NA | 2 | Gibert; Magda; Hazard | 10.1371/journal.pone.0139919 | 2015 |
| *Fritillaria biflora* | Liliaceae | NA | 2 | Yonezawa; Kinoshita; Watano; Zentoh | 10.1111/j.0014-3820.2000.tb01244.x | 2000 |
| *Gardenia actinocarpa* | Rubiaceae | NA | 2 | Osunkoya | 10.1016/S0006-3207(02)00417-2 | 2003 |
| *Gaura neomexicana coloradensis* | Onagraceae | NA | 9 | Floyd; Ranker | 10.1086/297607 | 1998 |
| *Gentiana pneumonanthe* | Gentianaceae | Marsh gentian | 1 | Oostermeijer; Brugman; de Boer; den Nijs | 10.2307/2261351 | 1996 |

| | | | | | | |
|---|---|---|---|---|---|---|
| *Gentianella campestris* | Gentianaceae | NA | 1 | Lennartsson; Oostermeijer | 10.1046/j.1365-2745.2001.00566.x | 2001 |
| *Geonoma pohliana weddelliana* | Arecaceae | NA | 1 | Souza; Martins | 10.1111/j.1442-9993.2006.01650.x | 2006 |
| *Geonoma schottiana* | Arecaceae | NA | 1 | Sampaio; Scariot | 10.1017/S0266467409990599 | 2010 |
| *Geranium sylvaticum* | Geraniaceae | NA | 1 | Ramula; Toivonen; Mutikainen | 10.1086/512040 | 2007 |
| *Geum rivale* | Rosaceae | NA | 2 | Kiviniemi | 10.1023/A:1015506019670 | 2002 |
| *Guarianthe aurantiaca* | Orchidaceae | NA | 3 | Mondragón | 10.1111/j.1442-1984.2009.00230.x | 2009 |
| *Helenium virginicum* | Compositae | NA | 1 | Adams; Marsh; Knox | 10.1016/j.biocon.2005.02.001 | 2005 |
| *Helianthemum juliae* | Cistaceae | NA | 1 | Marrero-Gómez; Oostermeijer; Carqué-Álamo; Bañares-Baudet | 10.1016/j.biocon.2007.01.010 | 2007 |
| *Helianthemum polygonoides* | Cistaceae | NA | 1 | Iriondo; Albert; Gimenez; Lozano; Escudero | 978-84-8014-746-0 | 2009 |
| *Helianthus divaricatus* | Compositae | Divaricate sunflower | 4 | Nantel; Gagnon | 10.1046/j.1365-2745.1999.00388.x | 1999 |
| *Heliconia acuminata* | Heliconiaceae | NA | 5 | Bruna | 10.1890/0012-9658(2003)084[0932:APPIFH]2.0.CO;2 | 2003 |
| *Heliconia metallica* | Heliconiaceae | NA | 4 | Schleuning; Huamán; Matthies | 10.1111/j.1365-2745.2008.01416.x | 2008 |
| *Heteropsis flexuosa* | Araceae | NA | 2 | Balcazar | 978-90-9027402-7 | 2013 |
| *Heteropsis macrophylla* | Araceae | NA | 2 | Balcazar | 978-90-9027402-7 | 2013 |
| *Heteropsis oblongifolia* | Araceae | NA | 2 | Balcazar | 978-90-9027402-7 | 2013 |
| *Hilaria mutica* | Poaceae | Tussock grass | 3 | Vega; Montaña | 10.1023/B:VEGE.0000048094.21994.74 | 2004 |

| | | | | | | |
|---|---|---|---|---|---|---|
| *Himatanthus drasticus* | Apocynaceae | Plumel | 1 | Baldauf; Correa; Ferreira; Santos | 10.1016/j.foreco.2015.06.022 | 2015 |
| *Horkelia congesta* | Rosaceae | Shaggy horkelia | 1 | Kaye; Benfield | NA | 2004 |
| *Hyparrhenia diplandra* | Poaceae | Tussock grass | 3 | Garnier; Dajoz | 10.1890/0012-9658(2001)082[1720:ESOALV]2.0.CO;2 | 2001 |
| *Hypericum cumulicola* | Hypericaceae | NA | 13 | Quintana-Ascencio; Menges; Weekley | 10.1046/j.1523-1739.2003.01431.x | 2003 |
| *Iris germanica* | Iridaceae | German Iris | 1 | Burns; Pardini; Schutzenhofer; Chung; Seidler; Knight | 10.1890/12-1310.1 | 2013 |
| *Jacquiniella leucomelana* | Orchidaceae | NA | 4 | Winkler; Hülber; Hietz | 10.1093/aob/mcp188 | 2009 |
| *Jacquiniella teretifolia* | Orchidaceae | NA | 3 | Winkler; Hülber; Hietz | 10.1093/aob/mcp188 | 2009 |
| *Juniperus procera* | Cupressaceae | NA | 1 | Couralet; Sass-Klaassen; Sterck; Bekele; Zuidema | 10.1016/j.foreco.2005.05.065 | 2005 |
| *Khaya senegalensis* | Meliaceae | African mabogany | 6 | Gaoue; Ticktin | 10.1111/j.1523-1739.2009.01345.x | 2010 |
| *Kosteletzkya pentacarpos* | Malvaceae | NA | 1 | Pino; Picó; Roa | 10.1111/j.1095-8339.2007.00628.x | 2007 |
| *Kummerowia striata* | Legumiosae | NA | 1 | Levin; Crandall; Knight | 10.1002/ecy.2681 | 2019 |
| *Laccosperma secundiflorum* | Arecaceae | NA | 1 | Kouassi; Barot; Gignoux; Bi | 10.1017/S0266467408005312 | 2008 |
| *Laserpitium longiradium* | Apiaceae | NA | 1 | Iriondo; Albert; Giménez; Lozano; Escudero | 978-84-8014-746-0 | 2009 |
| *Lechea cernua* | Cistaceae | NA | 2 | Maliakal Witt | NA | 2004 |
| *Lechea deckertii* | Cistaceae | NA | 3 | Maliakal Witt | NA | 2004 |
| *Leontopodium nivale alpinum* | Compositae | NA | 2 | Keller; Vittoz | 10.1007/s00035-014-0142-y | 2015 |

| | | | | | | |
|---|---|---|---|---|---|---|
| *Lepanthes eltoroensis* | Orchidaceae | NA | 4 | Tremblay; Ackerman | 10.1006/bijl.2000.0485 | 2001 |
| *Lepanthes rubripetala* | Orchidaceae | NA | 14 | Schödelbauerová; Tremblay; Kindlmann, Tremblay; Ackerman | 10.1007/s10531-009-9724-1, 10.1006/bijl.2000.0485 | 2010, 2001 |
| *Lepanthes rupestris* | Orchidaceae | NA | 14 | Tremblay; Ackerman, Tremblay; McCarthy | 10.1006/bijl.2000.0485, 10.1371/journal.pone.0102859 | 2001, 2014 |
| *Lepidium davisii* | Brassicaceae | Davis' peppergrass | 8 | Bernatus | NA | 1995 |
| *Liatris scariosa* | Compositae | Savanno Blazing Star | 4 | Ellis | 10.1890/11-1052.1 | 2012 |
| *Limonium carolinianum* | Plumbaginaceae | Salt marsh plant | 1 | Baltzer; Reekie; Hewlin; Taylor; Boates | 10.1139/b02-070 | 2002 |
| *Limonium delicatulum* | Plumbaginaceae | NA | 1 | Hegazy | 10.2307/2404462 | 1992 |
| *Limonium erectum* | Plumbaginaceae | NA | 1 | Iriondo; Albert; Giménez; Lozano; Escudero | 978-84-8014-746-0 | 2009 |
| *Limonium malacitanum* | Plumbaginaceae | NA | 1 | Iriondo; Albert; Giménez; Lozano; Escudero | 978-84-8014-746-0 | 2009 |
| *Lomatium bradshawii* | Apiaceae | NA | 2 | Kaye; Pendergrass; Finley; Kauffman | 10.1890/1051-0761(2001)011[1366:TEOFOT]2.0.CO;2 | 2001 |
| *Lomatium cookii* | Apiaceae | NA | 3 | Kaye; Pyke | 10.1890/0012-9658(2003)084[1464:TEOSTO]2.0.CO;2 | 2003 |
| *Lotus arinagensis* | Leguminosae | NA | 1 | Iriondo; Albert; Giménez; Lozano; Escudero | 978-84-8014-746-0 | 2009 |
| *Lupinus tidestromii* | Leguminosae | NA | 3 | Dangremond; Knight | 10.1890/09-0418.1 | 2010 |
| *Lycaste aromatica* | Orchidaceae | NA | 4 | Winkler; Hülber; Hietz | 10.1093/aob/mcp188 | 2009 |

| | | | | | | |
|---|---|---|---|---|---|---|
| *Machaerium cuspidatum* | Leguminosae | NA | 3 | Nabe-Nielsen | 10.1017/S0266467404001609 | 2004 |
| *Magnolia macrophylla dealbata* | Magnoliaceae | NA | 1 | Sánchez-Velásquez; Pineda-López | 10.1007/s10144-009-0161-5 | 2010 |
| *Mammillaria dixanthocentron* | Cactaceae | NA | 1 | Ramos Lopez | NA | 2007 |
| *Mammillaria hernandezii* | Cactaceae | NA | 2 | Rodriguez Ortega | NA | 2008 |
| *Mammillaria huitzilopochtli* | Cactaceae | NA | 2 | Flores Martinez, Flores-Martinez; Manzanero-Medino; Golubov; Montaña; Mandujano | NA, 10.1007/s11258-010-9737-6 | 2010 |
| *Mammillaria magnimamma* | Cactaceae | NA | 3 | Valverde; Quijas; Lopez-Villavicencio; Castillo | 10.1023/B:VEGE.0000021662.78634.de | 2004 |
| *Mammillaria napina* | Cactaceae | NA | 1 | Rodriguez Ortega | NA | 2008 |
| *Mammillaria solisioides* | Cactaceae | NA | 1 | Rodriguez Ortega | NA | 2008 |
| *Manilkara zapota* | Sapotaceae | NA | 1 | Cruz-Rodriguez; López-Villavicencio; Valverde | 10.1017/S0266467408005713 | 2009 |
| *Mauritia flexuosa* | Arecaceae | Canangucho; Morete | 1 | Holm; Miller; Cropper | 10.1111/j.1744-7429.2008.00412.x | 2008 |
| *Miconia albicans* | Melastomataceae | NA | 1 | Hoffmann | 10.2307/177080 | 1999 |
| *Mimulus cardinalis* | Phrymaceae | NA | 4 | Angert | 10.1890/0012-9658(2006)87[2014:DOCAMP]2.0.CO;2 | 2006 |
| *Mimulus lewisii* | Phrymaceae | NA | 3 | Angert | 10.1890/0012-9658(2006)87[2014:DOCAMP]2.0.CO;2 | 2006 |
| *Minuartia obtusiloba* | Caryophyllaceae | NA | 1 | Forbis; Doak | 10.3732/ajb.91.7.1147 | 2004 |

| | | | | | | |
|---|---|---|---|---|---|---|
| *Molinia caerulea* | Poaceae | Purple moor grass | 4 | Jacquemyn; Brys; Neubert | 10.1890/04-1762 | 2005 |
| *Myrsine guianensis* | Primulaceae | NA | 1 | Hoffmann | 10.2307/177080 | 1999 |
| *Nardostachys jatamansi* | Caprifoliaceae | NA | 1 | Ghimire; Gimenez; Pradel; McKey; Aumeeruddy-Thomas | 10.1111/j.1365-2664.2007.01375.x | 2008 |
| *Neobuxbaumia polylopha* | Cactaceae | NA | 1 | Arroyo-Cosultchi; Golubov; Mandujano | 10.1016/j.actao.2016.01.006 | 2016 |
| *Neotinea ustulata* | Orchidaceae | Burnt orchid | 1 | Shefferson; Tali | 10.1111/j.1365-2745.2006.01195.x | 2007 |
| *Oenothera deltoides* | Onagraceae | Antioch Dunes evening primrose | 4 | Thomson | 10.1111/j.1523-1739.2005.004108.x | 2005 |
| *Opuntia macrocentra* | Cactaceae | Purple prickly pear | 2 | Mandujano; Golubov; Huenneke | 10.1007/s10144-006-0032-2 | 2007 |
| *Opuntia microdasys* | Cactaceae | NA | 4 | Carrillo Angeles | NA | 2011 |
| *Orchis purpurea* | Orchidaceae | Lady orchid | 7 | Jacquemyns; Brys; Jongejans | 10.1890/08-2321.1 | 2010 |
| *Oxalis acetosella* | Oxalidaceae | NA | 3 | Berg | 10.1034/j.1600-0587.2002.250211.x | 2002 |
| *Oxytropis jabalambrensis* | Leguminosae | NA | 2 | Iriondo; Albert; Giménez; Lozano; Escudero | 978-84-8014-746-0 | 2009 |
| *Pachycereus pecten-aboriginum* | Cactaceae | NA | 3 | Morales-Romero; Godinez-Alvarez; Campo-Alves; Molino-Freaner | 10.1016/j.jaridenv.2011.09.005 | 2012 |
| *Paeonia officinalis* | Paeoniaceae | NA | 3 | Andrieu; Freville; Besnord; Vaudey; Gauthier; Thompson; Debussche | 10.1007/s10144-012-0346-1 | 2013 |
| *Panax quinquefolius* | Araliaceae | American ginseng | 1 | Shahi | NA | 2007 |

| | | | | | | |
|---|---|---|---|---|---|---|
| *Parkinsonia aculeata* | Leguminosae | NA | 1 | Raghu; Wilson; Dhileepan | 10.1111/j.1440-6055.2006.00556.x | 2006 |
| *Parolinia glabriuscula* | Brassicaceae | NA | 1 | Iriondo; Albert; Gimenez; Lozano; Escudero | 978-84-8014-746-0 | 2009 |
| *Paronychia pulvinata* | Caryophyllaceae | NA | 1 | Forbis; Doak | 10.3732/ajb.91.7.1147 | 2004 |
| *Periandra mediterranea* | Leguminosae | NA | 1 | Hoffmann; Solbrig | 10.1016/S0378-1127(02)00566-2 | 2003 |
| *Persoonia bargoensis* | Proteaceae | NA | 2 | McKenna | NA | 2007 |
| *Persoonia glaucescens* | Proteaceae | NA | 1 | McKenna | NA | 2007 |
| *Phyllanthus emblica* | Phyllanthaceae | Amla | 2 | Ellis; Williams; Lesica; Bell; Bierzychudek; Bowles; Crone; Doak; Ehrlén; Ellis-Adam; McEachern; Ganesan; Latham; Luijten; Kaye; Knight; Menges; Morris; Den Nijs; Oostermeijer; Quintana-Ascencio; Shelly; Stanley; Thorpe; Ticktin; Valverde; Weekley, Ticktin; Ganesan; Paramesha; Setty, | 10.1890/11-1052.1, 10.1111/j.1365-2664.2012.02156.x | 2012 |
| *Phyllanthus indofischeri* | Phyllanthaceae | Amla | 1 | Ticktin; Ganesan; Paramesha; Setty | 10.1111/j.1365-2664.2012.02156.x | 2012 |
| *Pinus lambertiana* | Pinaceae | NA | 1 | van Mantgem; Stephenson | 10.1111/j.1365-2745.2005.01007.x | 2005 |
| *Pinus maximartinezii* | Pinaceae | Blue pine; Mexican maxipinon | 1 | López-Mata | 10.1016/j.actao.2013.02.010 | 2013 |

| | | | | | | |
|---|---|---|---|---|---|---|
| *Pinus nigra* | Pinaceae | Black pine | 2 | Buckley; Brockerhoff; Langer; Ledgard; North; Rees | 10.1111/j.1365-2664.2005.01100.x | 2005 |
| *Pinus strobus* | Pinaceae | Eastern white pine | 3 | Münzbergová; Hadincová; Wild; Kindlmannová | 10.1371/journal.pone.0056953 | 2013 |
| *Pityopsis aspera aspera* | Compositae | Pineland silkgrass | 1 | Gornish | 10.1093/aobpla/plt041 | 2013 |
| *Plantago coronopus* | Plantaginaceae | NA | 7 | Villellas; Ehrlén; Olesen; Braza; García | 10.1111/j.1600-0587.2012.07425.x | 2013 |
| *Polemonium van-bruntiae* | Polemoniaceae | Jacob's ladder | 4 | Bermingham | 10.1007/s11258-010-9762-5 | 2010 |
| *Polygonella basiramia* | Polygonaceae | NA | 2 | Maliakal Witt | NA | 2004 |
| *Primula elatior* | Primulaceae | Oxlip | 7 | Jacquemyn; Brys | 10.1890/07-1908.1 | 2008 |
| *Primula veris* | Primulaceae | Cowslip | 3 | Endels; Jacquemyn; Brys; Hermy, Ehrlén; Syrjänen; Leimu; Garcia; Lehtilä | 10.1007/s11258-004-0026-0, 10.1111/j.1365-2664.2005.01015.x | 2005 |
| *Primula vulgaris* | Primulaceae | English primrose | 20 | Valdes; Garcia; Garcia; Ehrlen, Valverde; Silvertown | 10.1111/j.1600-0587.2013.00216.x, 10.1046/j.1365-2745.1998.00280.x | 2013, 1998 |
| *Prosopis glandulosa* | Leguminosae | Honey mesquite | 1 | Golubov; Mandujano; Franco; Montaña; Eguiarte; Lopez-Portillo | 10.1046/j.1365-2745.1999.00420.x | 1999 |
| *Prosopis laevigata* | Leguminosae | Smooth Mesquite | 1 | Bernal | NA | 2004 |
| *Prunus africana* | Rosaceae | Red stinkwood | 1 | Stewart | NA | 2001 |
| *Prunus serotina* | Rosaceae | Black cherry; Wild black cherry; Rum cherry | 2 | Sebert-Cuvillier; Paccaut; Chabrerie; Endels; Goubet; Decoq | 10.1016/j.ecolmodel.2006.09.005 | 2007 |

| | | | | | | |
|---|---|---|---|---|---|---|
| *Pseudomisopates rivas-martinezii* | Scrophulariaceae | NA | 2 | Iriondo; Albert; Giménez; Lozano; Escudero | 978-84-8014-746-0 | 2009 |
| *Pseudomitrocereus fulviceps* | Cactaceae | NA | 1 | Vite Gonzalez; Zavala Hurtado | NA | 1998 |
| *Pseudophoenix sargentii* | Arecaceae | Buccaneer palm; Sargent's cherry palm; Cherry palm | 3 | Durán; Franco, Maschinski; Duquesnel | NA, 10.1016/j.biocon.2006.07.012 | 1992, 2006 |
| *Pterocereus gaumeri* | Cactaceae | NA | 2 | Méndez; Duran; Olmsted | 10.1646/1601 | 2004 |
| *Ptychosperma macarthurii* | Arecaceae | NA | 1 | Liddle; Brook; Matthews; Taylor; Caley | 10.1016/j.biocon.2006.04.028 | 2006 |
| *Purshia subintegra* | Rosaceae | Arizona cliffrose | 3 | Maschinski; Baggs; Quintana-Ascencio; Menges | 10.1111/j.1523-1739.2006.00272.x | 2006 |
| *Pyrrocoma radiata* | Compositae | NA | 10 | Kaye; Pyke, Pfingsten | 10.1890/0012-9658(2003)084[1464:TEOSTO]2.0.CO;2, NA | 2003, 2013 |
| *Ramonda myconi* | Gesneriaceae | Pyrenean violet; Rosette mullein | 3 | Picó; Riba | 10.1023/A:1020310609348 | 2002 |
| *Rhododendron ponticum* | Ericaceae | NA | 8 | Salguero-Gómez | NA | 2004 |
| *Rosmarinus tomentosus* | Lamiaceae | NA | 1 | Iriondo; Albert; Giménez; Lozano; Escudero | 978-84-8014-746-0 | 2009 |
| *Rubus praecox* | Rosaceae | Himalayan blackberry | 3 | Lambrecht-McDowell; Radosevich | 10.1007/s10530-004-0870-9 | 2005 |
| *Rubus saxatilis* | Rosaceae | NA | 3 | Eriksson | 10.1007/BF02348412 | 1994 |
| *Rubus ursinus* | Rosaceae | Trailing blackberry | 3 | Lambrecht-McDowell; Radosevich | 10.1007/s10530-004-0870-9 | 2005 |

| | | | | | | |
|---|---|---|---|---|---|---|
| *Rumex rupestris* | Polygonaceae | NA | 1 | Iriondo; Albert; Giménez; Lozano; Escudero | 978-84-8014-746-0 | 2009 |
| *Sabal yapa* | Arecaceae | NA | 2 | Pulido; Valverde; Caballero | 10.1017/S0266467406003877 | 2007 |
| *Santolina melidensis* | Compositae | NA | 1 | Iriondo; Albert; Giménez; Lozano; Escudero | 978-84-8014-746-0 | 2009 |
| *Sapium sebiferum* | Euphorbiaceae | *Triadica sebifera*; Chinese tallow tree; Florida aspen; Chicken tree; Gray popcorn tree; Candleberry tree | 4 | Renne | NA | 2001 |
| *Saponaria bellidifolia* | Caryophyllaceae | NA | 4 | Csergő; Molnár; Garcia | 10.1007/s10144-010-0249-y | 2011 |
| *Sarcocapnos baetica* | Papaveraceae | NA | 1 | Salinas; Suárez; Blanca | 10.1139/b02-013 | 2002 |
| *Sarcocapnos enneaphylla* | Papaveraceae | NA | 1 | Salinas; Suárez; Blanca | 10.1139/b02-013 | 2002 |
| *Sarcocapnos pulcherrima* | Papaveraceae | NA | 1 | Salinas; Suárez; Blanca | 10.1139/b02-013 | 2002 |
| *Sarracenia purpurea* | Sarraceniaceae | Northern pitcher plant | 3 | Gotelli; Ellison, Tendland | 10.1890/04-0479, NA | 2006, 2011 |
| *Saussurea medusa* | Compositae | Snow Lotus | 1 | Law; Salick; Knight | 10.1007/s11258-010-9761-6 | 2010 |
| *Saxifraga aizoides* | Saxifragaceae | NA | 3 | Marcante; Winkler; Erschbamer | 10.1093/aob/mcp047 | 2009 |
| *Saxifraga cotyledon* | Saxifragaceae | NA | 3 | Dinnetz; Nilsson | 10.1023/A:1015593311183 | 2002 |
| *Saxifraga tridactylites* | Saxifragaceae | NA | 4 | Dostál | 10.1111/j.1654-1103.2007.tb02519.x | 2007 |

| | | | | | | |
|---|---|---|---|---|---|---|
| *Scaphium macropodum* | Malvaceae | NA | 4 | Yamada; Zuidema; Itoh; et al. | 10.1111/j.1365-2745.2006.01209.x | 2007 |
| *Senecio filaginoides* | Compositae | NA | 1 | Cipriotti; Aguiar | 10.1111/j.1654-109X.2011.01138.x | 2012 |
| *Shorea acuminata* | Dipterocarpaceae | NA | 3 | Yamada; Yamada; Okuda; Fletcher | 10.1007/s00442-012-2529-z | 2013 |
| *Shorea bracteolata* | Dipterocarpaceae | NA | 3 | Yamada; Yamada; Okuda; Fletcher | 10.1007/s00442-012-2529-z | 2013 |
| *Shorea leprosula* | Dipterocarpaceae | Meranti Tembaga | 4 | Visser; Jongejans; van Breugel; Zuidema; Chen; Kassim; de Kroon, Yamada; Yamada; Okuda; Fletcher | 10.1111/j.1365-2745.2011.01825.x, 10.1007/s00442-012-2529-z | 2011, 2013 |
| *Shorea maxwelliana* | Dipterocarpaceae | NA | 3 | Yamada; Yamada; Okuda; Fletcher | 10.1007/s00442-012-2529-z | 2013 |
| *Shorea ovalis* | Dipterocarpaceae | NA | 3 | Yamada; Yamada; Okuda; Fletcher | 10.1007/s00442-012-2529-z | 2013 |
| *Silene acaulis* | Caryophyllaceae | Cushion plant | 5 | Morris; Doak | 10.2307/2446413 | 1998 |
| *Sporobolus heterolepis* | Poaceae | Prairie dropseed | 1 | Dalgleish; Kula; Hartnett; Sandercock | 10.3732/ajb.2007277 | 2008 |
| *Stenocactus crispatus* | Cactaceae | NA | 1 | Lopez Flores; Navarro Carbajal | NA | 2009 |
| *Stenocereus eruca* | Cactaceae | NA | 1 | Clark-Tapia | NA | 2004 |
| *Stipa calamagrostis* | Poaceae | NA | 1 | Guardia; Raventos; Caswell | 10.1046/j.1365-2745.2000.00504.x | 2000 |
| *Succisa pratensis* | Caprifoliaceae | NA | 6 | Wallin; Svensson, Milden | 10.1007/s12224-012-9123-3, NA | 2012, 2005 |
| *Swietenia macrophylla* | Meliaceae | Big leaf mahogany | 1 | Verwer; Peña-Claros; van der Staak; Ohlson-Kiehn; Sterck | 10.1111/j.1365-2664.2008.01564.x | 2008 |
| *Syzygium jambos* | Myrtaceae | NA | 1 | Brown; Spector; Wu | 10.1111/j.1365-2664.2008.01550.x | 2008 |

| | | | | | | |
|---|---|---|---|---|---|---|
| *Tetramolopium arenarium* | Composita e | NA | 1 | Aplet; Laven; Shaw | NA | 1994 |
| *Tetraneuris herbacea* | Composita e | Eastern four-nerved daisy, Stemless four-nerved daisy | 3 | Campbell; Husband | 10.1038/sj.hdy.6800653 | 2005 |
| *Thlaspi perfoliatum* | Brassicac eae | NA | 1 | Burns; Pardini; Schutzenhofer; Chung; Seidler; Knight | 10.1890/12-1310.1 | 2013 |
| *Tillandsia brachycaulos* | Bromeliac eae | NA | 1 | Mondragón; Dúran; Ramírez; Valverde | 10.1017/S0266467403001287 | 2004 |
| *Tillandsia deppeana* | Bromeliac eae | NA | 1 | Winkler; Hülber; Hietz | 10.1016/j.baae.2006.05.003 | 2007 |
| *Tillandsia juncea* | Bromeliac eae | NA | 1 | Winkler; Hülber; Hietz | 10.1016/j.baae.2006.05.003 | 2007 |
| *Tillandsia macdougallii* | Bromeliac eae | NA | 1 | Mondragón; Ticktin | 10.1111/j.1523-1739.2011.01691.x | 2011 |
| *Tillandsia multicaulis* | Bromeliac eae | NA | 1 | Winkler; Hülber; Hietz | 10.1016/j.baae.2006.05.003 | 2007 |
| *Tillandsia violacea* | Bromeliac eae | NA | 1 | Mondragón; Ticktin | 10.1111/j.1523-1739.2011.01691.x | 2011 |
| *Tolumnia variegata* | Orchidace ae | NA | 1 | Calvo | 10.2307/1940473 | 1993 |
| *Trillium grandiflorum* | Melanthia ceae | NA | 13 | Knight | 10.3732/ajb.90.8.1207 | 2003 |
| *Trillium ovatum* | Melanthia ceae | Western trillium | 1 | Ream | NA | 2011 |
| *Trillium persistens* | Melanthia ceae | Persistent trillium | 5 | Plank | NA | 2010 |
| *Trollius europaeus* | Ranuncula ceae | NA | 3 | Lemke; Salguero-Gomez | 10.1007/s10144-015-0519-9 | 2015 |
| *Trollius laxus* | Ranuncula ceae | Globeflower | 1 | Scanga; Leopold | 10.1016/j.biocon.2012.01.061 | 2012 |
| *Tsuga canadensis* | Pinaceae | Eastern hemlock | 2 | Lamar; McGraw | 10.1016/j.foreco.2005.02.056 | 2005 |

| | | | | | | |
|---|---|---|---|---|---|---|
| *Vella pseudocytisus* | Brassicaceae | NA | 6 | Iriondo; Albert; Giménez; Lozano; Escudero, Dominguez Lozano; Moreno Saiz; Schwartz | 978-84-8014-746-0, 10.1016/j.jnc.2011.05.005 | 2009, 2011 |
| *Verbascum fontqueri* | Scrophulariaceae | NA | 2 | Iriondo; Albert; Giménez; Lozano; Escudero | 978-84-8014-746-0 | 2009 |
| *Veronica arvensis* | Plantaginaceae | NA | 5 | Burns; Pardini; Schutzenhofer; Chung; Seidler; Knight, Dostal | 10.1890/12-1310.1, 10.1658/1100-9233(2007)18[91:PDOAIP]2.0.CO;2 | 2013, 2007 |
| *Verticordia staminosa* | Myrtaceae | NA | 1 | Yates; Ladd; Coates; McArthur | 10.1071/BT06032 | 2007 |
| *Viola elatior* | Violaceae | NA | 2 | Eckstein; Danihelka; Otte | 10.2478/s11756-009-0002-1 | 2009 |
| *Viola pumila* | Violaceae | NA | 2 | Eckstein; Danihelka; Otte | 10.2478/s11756-009-0002-1 | 2009 |
| *Vitaliana primuliflora* | Primulaceae | NA | 1 | Iriondo; Albert; Giménez; Lozano; Escudero | 978-84-8014-746-0 | 2009 |
| *Vriesea sanguinolenta* | Bromeliaceae | NA | 4 | Zotz | 10.1016/j.actao.2005.05.009 | 2005 |
| *Zamia inermis* | Zamiaceae | NA | 1 | Octavio-Aguilar; Rivera-Fernández; Iglesias-Andreu; Vovides; De Cáceres-González | 10.1007/s10531-016-1270-z | 2017 |
| *Zea diploperennis* | Poaceae | NA | 1 | Sanchez-Velazquez; Ezcurra; Martinez-Ramos; Alvarez-Buylla; Lorente | 10.1046/j.1365-2745.2002.00702.x | 2002 |

### Demographic traits calculation

From the selected MPMs, we extracted vital rates, life-history traits, and transient dynamics to study how they respond to different levels of human pressure. The mean unmanipulated MPM ***A*** has been divided into 3 submatrices ***U***, ***F*** and ***C.*** With ***A*** = ***U*** + ***F*** + ***C***. The submatrix ***U*** contains survival-dependent demographic processes, such as progression and retrogression from one stage to another; the submatrix ***F*** contains sexual per-capita contributions; the submatrix ***C*** contains clonal per-capita contributions as we chose to separate the two reproductive outputs: sexual and asexual.

Among the key traits to understand species' demographic strategies, vital rates allow to describe precisely how species allocate resources across the different life stages. Vital rates have the advantage of being easily extracted from the MPMs in a process that is described in this paragraph. Hence, to extract the progressive growth rate (later described as growth rate), the shrinkage rate (rate of retrogression to a less developed stage), the rate of sexual reproduction and the clonal rates of stage j, we summed the elements of the $j^{th}$ column corresponding to posterior stages of the matrix ***U***, the elements of the $j^{th}$ column corresponding to anterior stages of the matrix ***U***, all the elements of the $j^{th}$ column of the matrix ***F*** and all the elements of the $j^{th}$ column of the matrix ***C***. We then averaged the rates for each stage weighted by their contribution at the demographic equilibrium to get the rate at the population level. This contribution of each stage at the demographic equilibrium (i.e., the distribution of the different stages at the demographic equilibrium), is stated by the vector ***w***, which is the right eigenvector of ***A***[5]. This vector ***w*** represents the proportion of individuals in each stage when the population is at the equilibrium which corresponds to a situation where the growth rate is stable and here equal to 1 and the proportion does not change from one period to another. The survival rate of stage j is obtained by averaging the sum of all the columns of the matrix ***F*** weighted by their contribution at the demographic equilibrium. As an illustration to get the rate of sexual reproduction and the growth rate the formulas are respectively the following: $\sum_j (\sum_i f_{ij}) w_j$, and $\sum_j (\sum_{i>j} u_{ij}) w_j$ with $f_{ij}$ being the element of ***F*** in column j, row I and $u_{ij}$ being the element of ***U*** in column j, row i.

**Table S3: List of the demographic traits used in the analysis**. $\hat{A}$ being the standardized population model obtained by dividing A, the matrix population model, by the maximum eigenvalues $\lambda_{max}$. $l_x$ being the age-specific survival schedule, i.e, the survival to age/stage x. $m_x$ is the age-specific fertility schedule

| Trait ID | Trait name | Calculation | Description |
|---|---|---|---|
| **Life history traits** | | | |
| $R_0$ | Reproductive rate | $R_0 = \int_0^{\infty} l_x m_x dx$ | Average number of offspring produced by a female in a population in her lifetime |
| S | Degree of iteroparity | $S = -e^{-\log(\lambda_1)} l_x m_x \log(e^{-\log(\lambda_1)} l_x m_x)$ | Describe the range between iteroparous and semelparous strategies based on |

| | | | |
|---|---|---|---|
| | | | the Gini index S≈0 corresponds to highly semelparous species and S>>0 implies high degree of iteroparity. |
| T | Generation time | $T = \frac{\log{(R_0)}}{\log{(\lambda_1)}}$ | Average time required for an individual in the population to replace itself |
| $L_\alpha$ | Age at first reproduction | $L_\alpha$ | Mean age of an individual in the population at the first reproduction |
| H | Survivorship curve type | $H = \frac{-\log{(l_x)}l_x}{\sum l_x}$ | Shape of the age-specific survivorship curve lx as quantified by Keyfitz' entropy (H) H > 1, =1, <1 correspond to species whose mortality hazards increase, stay |

| | | | |
|---|---|---|---|
| | | | constant or decrease with age respectively |
| pRep | Probability of reproduction | $p_{rep}$ | Average probability for an individual in the population to reproduce and get offsprings |
| **Transient dynamics** | | | |
| ρ | Damping ratio | $\rho = \frac{\overline{\lambda_1}}{\lvert\overline{\lambda_2}\rvert}$ | Value describing how quickly the population return to a stable growth rate after a disturbance[5]. Damping ratio of 1 described a system that return to rest almost immediately without oscillation. |
| $P_i$ | Period of oscillation | $P = \frac{2\pi}{\arctan\left(\frac{\Im(\bar{\lambda}_2)}{\Re(\bar{\lambda}_2)}\right)}$ | Average expected time of one oscillation of the population growth rate after a disturbance[5] |

| | | | |
|---|---|---|---|
| | | Where $\Im(\bar{\lambda}_2)$ is the imaginary parts of $\bar{\lambda}_2$ and $\Re(\bar{\lambda}_2)$ its real part | |
| $\overline{\rho_1}$ | Reactivity | $\overline{\rho_1} = \lVert \hat{A} \rVert_1$ | Maximum population growth rate in a single timestep, relative to stable growth rate[4,6] |
| $\underline{\rho_1}$ | First step attenuation | $\underline{\rho_1} = minCS(\hat{A})$ | Minimum population growth in a single timestep, relative to stable growth rate[4,6] |

### Data global representation

Plotting the different localities of the 921 studied populations showcases their quasi-global distribution, though not exempt of important biogeographic biases (Fig. S1). Demographic data archived in COMPADRE and COMADRE can come from any species and location around the world. However, most of the studies are in North America and Europe. Asia, Oceania, and South America are currently poorly covered[7].

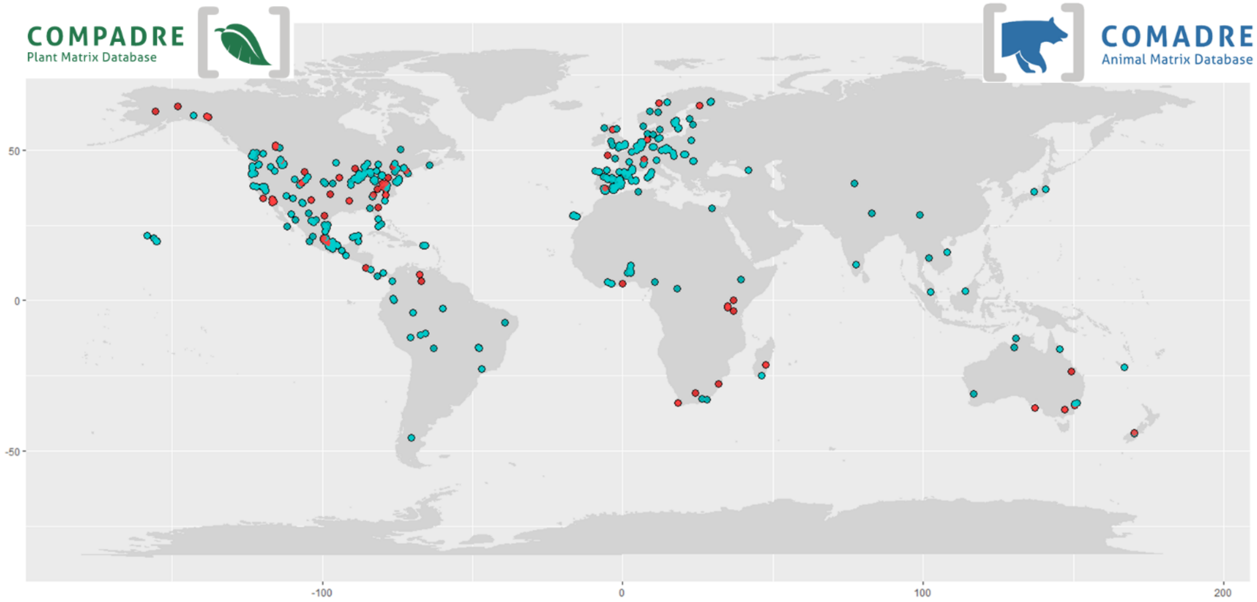

**Figure S1.** Map showing the global distribution of the natural populations used in this study. Light blue points represent plants, while light red ones represent animals. Importantly, some geographic areas are not well represented, such as Asia, Africa, or South America.

**Table S4. Representativity of the different kingdoms in our database**. For each kingdom we give the number of species (# species data) in our data, an approximation of the total estimated number of species in the realm (# species kingdom) and the corresponding coverage of the whole kingdom (% coverage). For number taken out of the paper *How Many Species Are There on Earth and in the Ocean?* from Mora et al., only the number of described species has been taken.

| Kingdom | # species data | # species kingdom | % coverage |
|---|---|---|---|
| Animalia | 45 | 1 900 000[8] | 0.002% |
| Plantae | 279 | 350 000[9] | 0.080% |
| Fungi | 1 | 44 000[10] | 0.002% |
| Algae* taken as Chromista here | 1 | 18 000[10] | 0.006% |

### Phylogenetic information gathering

Phylogenetic relatedness can generate biases in statistical analyses that neded to be consider. Dealing with several different species across the tree of life with sometimes several populations of the same species we had to consider the phylogenetic relatedness within our analyses. Therefore, we collected phylogenetic information at the species level to implement a series of phylogenetically corrected analyses. The phylogenetic tree of animals has been created with TimeTree[11] and the one for animals was generated using V.phylomaker[12]. For the animals' tree, three species were missing using TimeTree, so they were included by looking through the literature for the distance that separate them from their closest relatives in our tree. To do so, we looked in other phylogenetic trees such as the Open Tree Of Life[13,14].

We created a population level phylogeny to be able to correct for the phylogenetic proximity of the sampled population. To accommodate the fact that for some species we had more than one population (Table S1, Table S2), we modified the trees to include multiple populations within species. To do so, we added a new population on the tree branch of the species, each population being separated by a very small distance ($\varepsilon$= 0.0000001 normalized units), thus assuming that populations within a same species are very closely related. The distance ε was chosen as the smallest branch length that was detected by the programming software and that render a non polytomic tree. Then we did this step again, fitting the n+1 population at a distance of n*ε of the initial population in the phylogenetic tree. Thus, keeping a very small distance of ε between each species and avoiding the appearance of polytomies within the tree. For instance, for a third population, we fitted at a distance of 2ε of the initial population and hence a distance of ε of the second population that have been previously added on the tree. The order in which each population was affected to a branch tip was random.

Population level phylogenies were not affected by changes in the population structure. To make sure that the modification implemented in our phylogenetic trees to accommodate multiple populations per species did not affect the results of the analyses, we quantified the sensitivity of phylogenetic signals to changes in the order in which different populations within a given species diverged from each other. To do so, we randomly created 101 trees for plants and animals with random organisation of the populations within them. Then, we picked one of trees to be the reference tree and we compared the phylogenetic signal emerging from this tree architecture to the phylogenetic signals from the remaining 100 other trees. To quantify the phylogenetic signal, we used Pagel's *λ*[15], Blomberg's K , and Moran's I[17,18]. These metrics are all adapted to measure whether closely related species tend to resemble to each other, however, they do not rely all on the same assumptions. For instance, if they all test the null hypothesis of no phylogenetic signal, Moran's I test will compare the signal of the data with the signal obtained from random permutations of the tips while Pagel's λ is based on the comparison of signal obtained in the data and the signal obtained under the results of a hypothetical evolution under a Brownian motion. The phylogenetic signals we report in the main manuscript are insensitive to population permutations. Indeed, for Pagel's *λ* and Moran's I, no significant differences are found between the phylogenetic signals of all trees (Pagel's *λ*: $t_{99}$ = -2,224, $P < 0.05$; Moran's I: $t_{99}$ = -7.182, $P < 0.001$). However, for Blomberg's K, 30% of the trees produced an estimate that is significantly different from the baseline tree. This might be because the structure of the test itself, that relies on the permutation of the tip's values, is unbalanced by the fact that some species present multiple populations while other don't. To overcome this issue, we chose a reference tree that was rendering a Blomberg's K index that was unsensitive to permutations. Which, in a more complex language, is the phylogenetic tree that led us to accept the null of hypothesis of our *t*-test that stipulate a conformity of the sample values to the reference value, or in other word which say that whatever permutation we were doing in the tips order of the different populations, the Blomberg's K index of our reference tree was not statistically different of the one of the other trees generated.

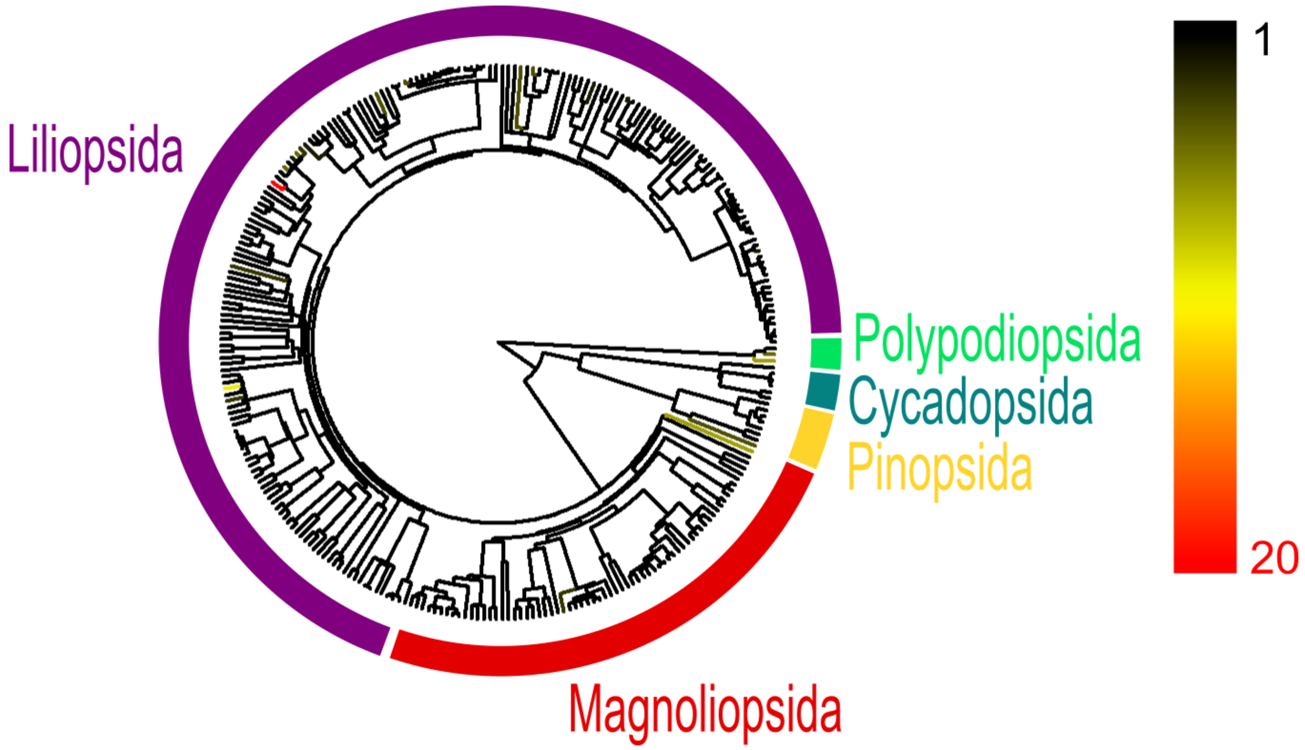

**Figure S2.** Resolved species-level phylogenetic tree of the 279 plant species used in our analyses. The phylogeny was obtained from V.Phylomaker[12] and modified to color the tip edges according to the number of different populations per species (Table S2). The taxonomic class to which the populations belong is shown around the tree. The legend shows the color gradient corresponding to the number of populations per species.

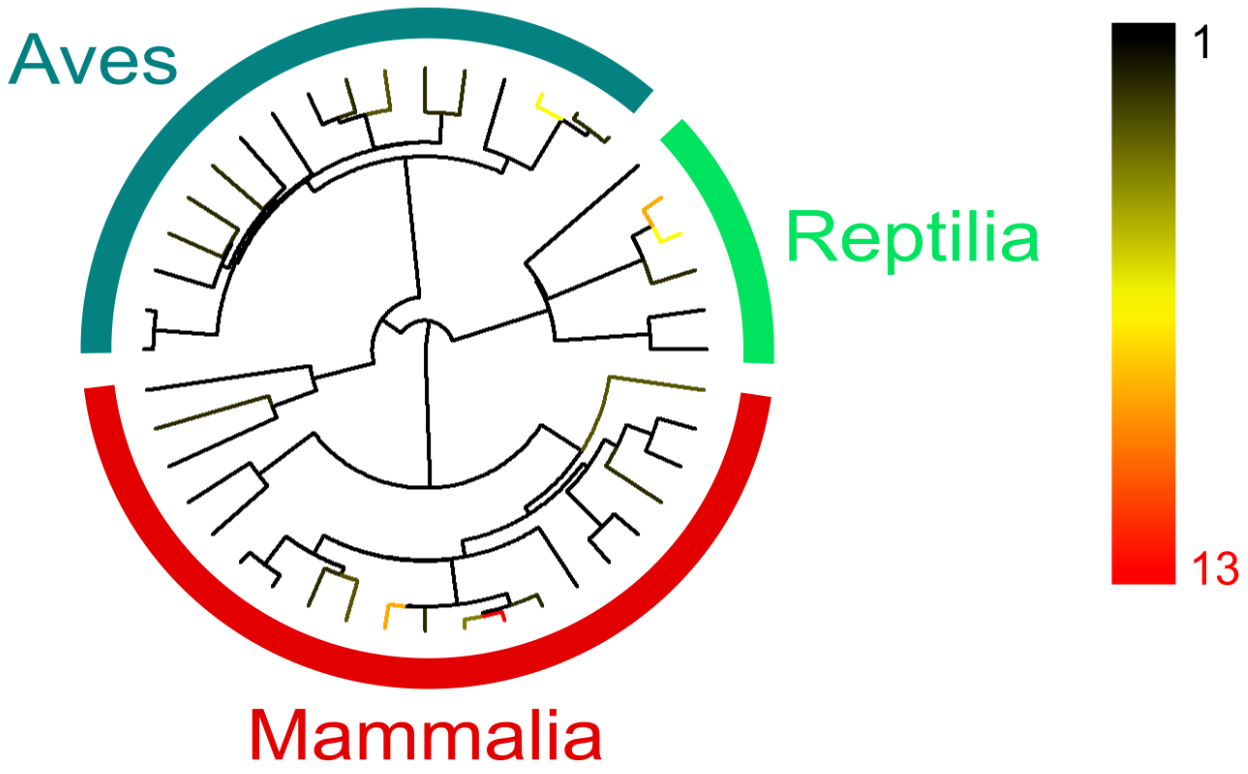

**Figure S3.** Resolved species-level phylogenetic tree of the 45 animal species used in our analyses. The phylogeny is issued from Timetree[11] and modified to color the species tips according to the number of populations per species (Table S1). The legend shows the color gradient corresponding to the number of populations per species.

### Linking traits and HFP

To identify the demographic traits that best explain the axes of human impact (Fig. 1), we ran a random tree forest analysis[19]. This analysis ensured that the posterior analyses (Table 1 and Tables S6, S7, S8, S9) avoided over-analysing the data. The results confirmed the importance of incorporating phylogenetic corrections, as certain taxonomic groups did tend to cluster together.

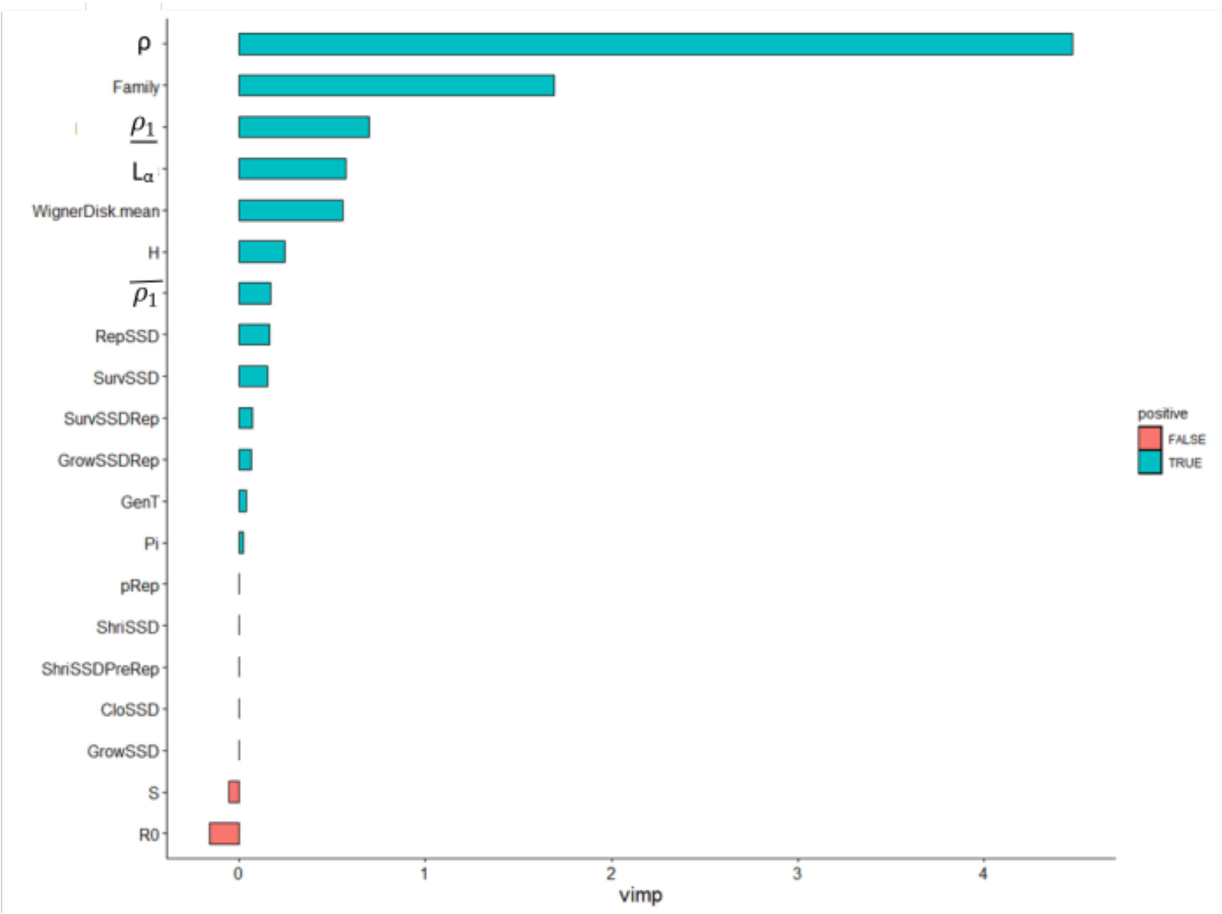

**Figure S4.** Results of the random forest analysis to study how animal demographic traits explain the first axis of the HFP-PCA (human presence). The total variance explained by all the traits is 39.07%. Blue bars represent positive correlations, while red ones negative correlations. The higher the vimp score is, the better the trait explains the PC1 (human presence; see Fig. 1).

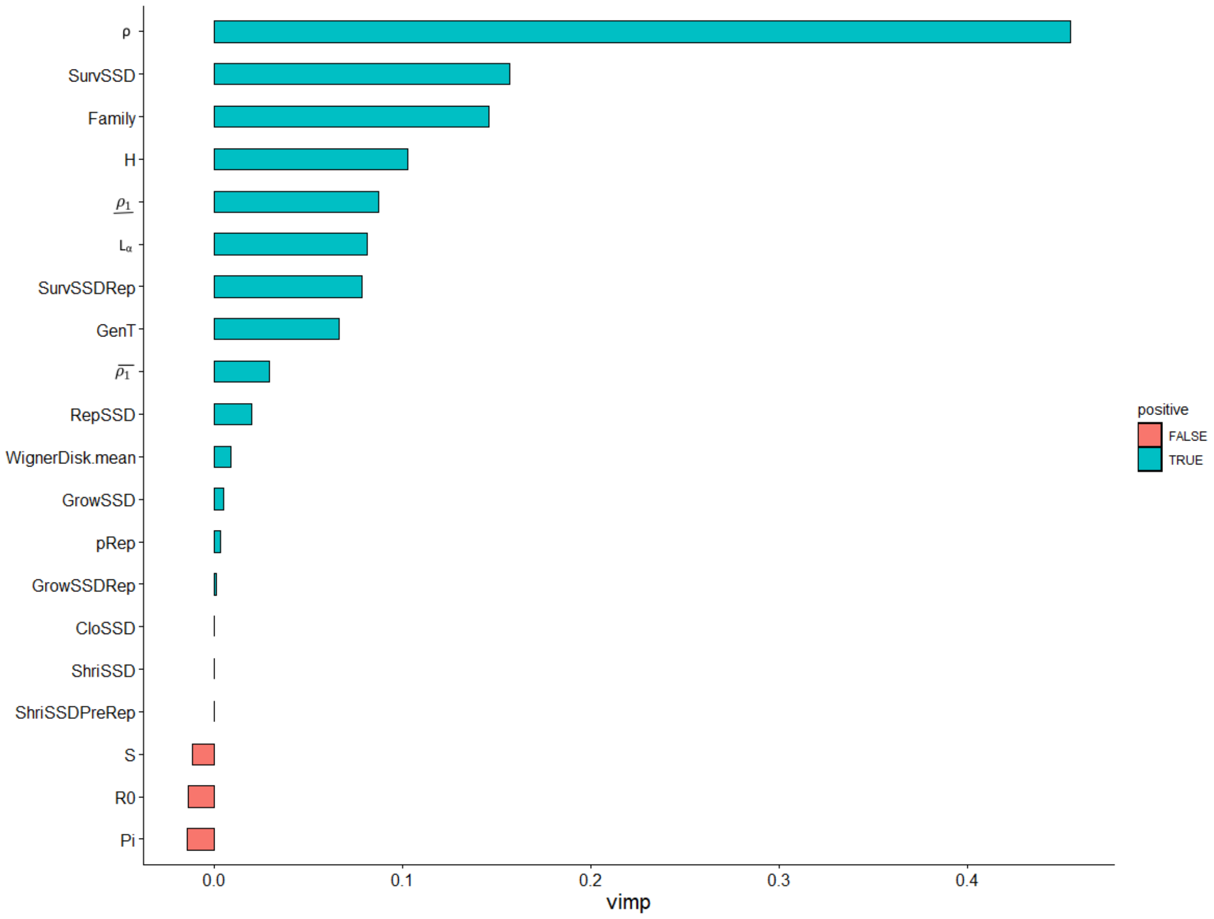

**Figure S5.**: Results of the random forest analysis to study how animal demographic traits explain the second axis of the HFP-PCA (agricultural land-use). The total variance explained by all the traits is 39.07%. Blue bars represent positive correlations, while red ones negative correlations. The higher the vimp score is, the better the trait explains the PC2 (agricultural land use; see Fig. 1).

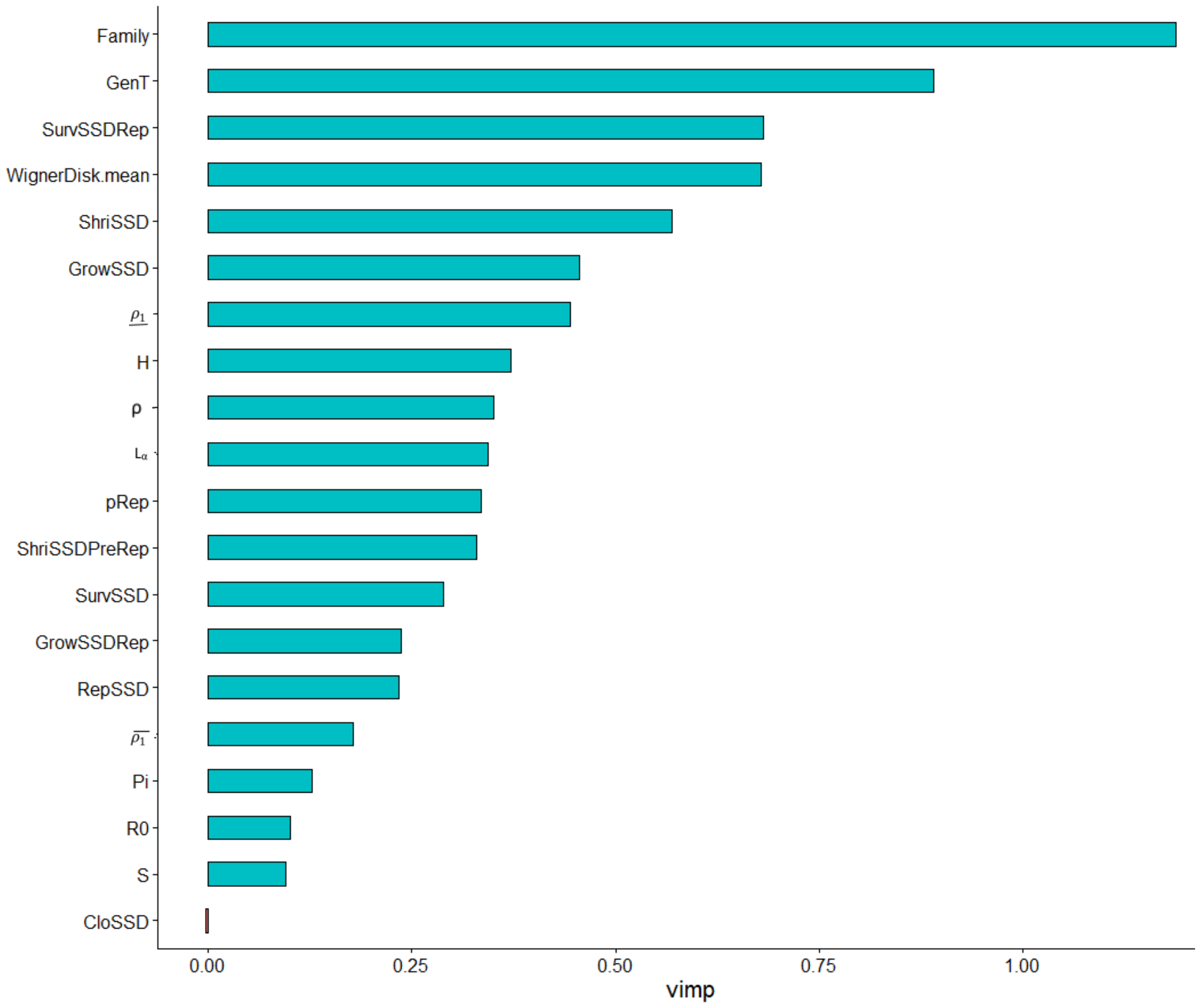

**Figure S6.** Results of the random forest analysis to study how plant demographic traits explain the first axis of the HFP-PCA (human presence). The total variance explained by all the traits is 35.49%. Blue line stands for positive correlation and red one for negative one. The higher the vimp score is, the better the trait explains the PCA-axis.

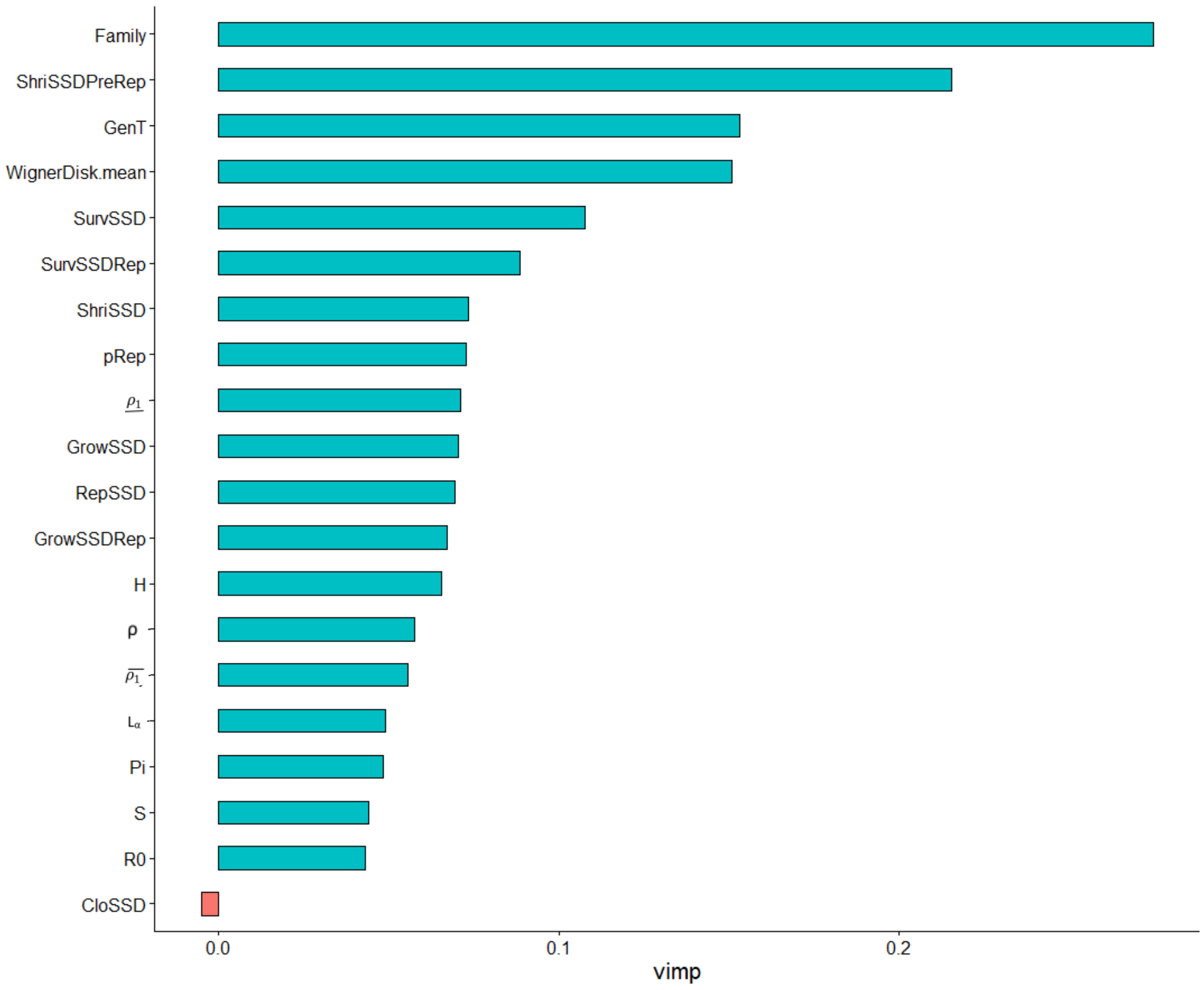
Family
ShriSSDPreRep
GenT
WignerDisk.mean
SurvSSD
SurvSSDRep
ShriSSD
pRep
$\underline{\rho_1}$
GrowSSD
RepSSD
GrowSSDRep
H
ρ
$\overline{\rho_1}$
$L_\alpha$
Pi
S
R0
CloSSD
0.0
0.1
0.2
vimp

**Figure S7.** Results of the random forest analysis to study how plant demographic traits explain the second axis of the HFP-PCA (agricultural land-use). Blue line stands for positive correlation and red one for negative one. The higher the vimp score is, the better the trait explains the PCA-axis.

### Appendex S2: Extended results

#### Human Footprint PCA

**Table S5.** Summary of the PCA with the 8 components of Human Footprint from Venter and collaborators. The eight components of Human footprint are: (1) human population density, (2) roads, (3) navigable waterways, (4) railways, (5) light pollutions, (6) built environment, (7) extensive pastures, and (8) intense croplands. The first column shows the eigenvalue of the PCA axes, the second column the percentage of variance explained. The figure with the axes selected can be found in the main corpus of the paper (Fig. 1).

| PCA - axis | Eigenvalue | Percentage of variance explained |
|---|---|---|
| 1$^{st}$ dimension | 2.869 | 35.862 |
| 2$^{nd}$ Dimension | 1.330 | 16.626 |
| 3$^{rd}$ Dimension | 1.098 | 13.723 |
| 4$^{th}$ Dimension | 0.963 | 12.038 |
| 5$^{th}$ Dimension | 0.681 | 8.516 |
| 6$^{th}$ Dimension | 0.562 | 7.029 |

| | | |
|---|---|---|
| 7th Dimension | 0.417 | 5.213 |
| 8th Dimension | 0.079 | 0.992 |

### Phylogenetic signal in resilience framework

**Table S6.** Phylogenetic signal analyses for the different aspects of the resilience frameworks. This table includes results for plants and animals. Results are given as the phylogenetic signal with the *P*-value between parentheses. Pagel's λ values are running between 0 and 1. Values close to 0 indicate no correlation between species, while values close to 1 indicate a strong phylogenetic signal (i.e., correlation between species equal to the Brownian expectation of evolution of traits).

| Phylogenetic signals plants and animals | |
|---|---|
| Trait | Pagel's *λ* |
| Resistance | 0.958 (*P* < 0.001)<br>0.936 (P < 0.001) |
| Speed of recovery | 0.970 (*P* < 0.001)<br>0.983 (P < 0.001) |
| Compensation | 0.989 (P < 0.001)<br>0.947 (P < 0.001) |

### Linear model in the resilience framework

Further, Table S6 shows the results of the linear model shown in the figure 3 and 4 of the main body of the paper. It shows that for plants the speed of recovery is significantly higher close to human presence while compensation is decreasing. However, close to farmlands plant species become more likely to experience sudden increase in population size after a disturbance (i.e., their compensation capacity increase). For animals, the mean resistance is significantly increasing along the axis of human presence, while it is decreasing when species are in areas with intensive agriculture. This trend appears also for animals with a short-distance mobility but not for those with a long-distance mobility. Ultimately, the compensation ability of species with a short-distance mobility is decreasing closer to farmlands.

**Table S7.** Results of the linear regression of the different components of the resilience framework along the axis of human pressure. This table includes the analyses for plants, animals, and subgroup of animals with low and high mobility ranges. Results are given in the following format: estimation of the effect ($t$-test$_{\text{degree of freedom}}$, $P$-value). Underlined $P$-values are statistically significant (*i.e.,* < 0.05).

| Results of linear regression for plants and animals | | |
|---|---|---|
| Trait | Human presence | Agricultural land use |
| Resistance | 0.017 ($t_{788}$ = 1.508, $P$ = 0.132)<br>0.037 ($t_{107}$ = 2.357, P = <u>0.020</u>) | -0.018 ($t_{788}$ = -0.889, $P$ = 0.374)<br>-0.120 ($t_{107}$ = -3.872, P < <u>0.001</u>) |
| Speed of recovery | 0.017 ($t_{788}$ = 2.976, $P$ = <u>0.003</u>)<br>-0.019 ($t_{107}$ = -0.526, P = 0.600) | 0.009 ($t_{788}$ = 0.860, $P$ = 0.390)<br>0.069 ($t_{107}$ = 0.917, P = 0.361) |
| Compensation | -0.041 ($t_{788}$ = -2.07, $P$ = <u>0.039</u>)<br>0.001 ($t_{107}$ = 0.064, P = 0.949) | 0.102 ($t_{788}$ = 2.924, $P$ = <u>0.004</u>)<br>-0.030 ($t_{107}$ = -0.952 P = 0.343) |
| Residual of linear regression for animals with long-distance mobility and short-distance mobility | | |
| Trait | Human presence | Agricultural land use |
| Resistance | 0.030 ($t_{71}$ = 1.954, $P$ = 0.055)<br>0.097 ($t_{34}$ = 2.510 P = <u>0.017</u>) | -0.076 ($t_{71}$ = -2.362, $P$ = <u>0.021</u>)<br>-0.287 ($t_{34}$ = -4.694, P < <u>0.001</u>) |
| Speed of recovery | -0.042 ($t_{71}$ = -0.856, $P$ = 0.395)<br>-0.019 ($t_{34}$ = -1.010, P = 0.319) | 0.106 ($t_{71}$ = 1.021, $P$ = 0.311)<br>0.064 ($t_{34}$ = 1.916 P = 0.064) |
| Compensation | -0.002 ($t_{71}$ = -0.191, $P$ = 0.849)<br>0.081 ($t_{34}$ = 1.971, P = 0.057) | -0.007 ($t_{71}$ = -0.418, $P$ = 0.678)<br>-0.178 ($t_{34}$ = -2.374, P = <u>0.023</u>) |

### Further demographic traits analyses

To study the distribution of demographic rates along gradients of human pressure, in addition of looking at classic linear models (see Table S4), we developed a new method incorporating both phylogenetic and spatial corrections as well as correction for body-size. However, to check that this new method produces reliable results, we also ran all these corrections separately and compared the different results. The “residual method” is the one presented in the paper, and that combined all the different corrections using the residuals of a regression of the demographic traits against size that accounted for spatial autocorrelation into the MCMCglmm. The other methods that run separately spatial and phylogenetic correction does not required this use of residuals and are hence, more reliable statistically. The model that incorporates spatial correction is a Generalized Additive Model (gam) with a tensor smooth taking into account the geographic coordinates of the populations, while the phylogenetically corrected method is a MCMCglmm that takes into account the variance related to the phylogenetic proximity.

The analysis of the results of the three different models produces congruent results (see Tables S6-S10). In summary, the residual methods render, overall, similar results as the two other methods that do not use residuals. Naturally, however, the residual method produces fewer significant results, as it accounts for more sources of autocorrelation. Also, models that just take spatial corrections into account tend to significantly explain more variation in traits values, as illustrated by generation time and survivorship in plants and probability of reproduction in animals.

We also ran all the different methods for sub selection of animal species to study the different mobility level (*i.e.* high *vs.* low mobility). Comparing species with a high mobility with species with a low mobility, it turns out that the two groups are not affected similarly by human activities. Hence, population with a lower mobility are likely to experience a shorter drop in population size after a disturbance (First Step Attenuation) in dense agricultural areas compared to populations in pristine habitat. Consequently, low mobile animal populations living in agricultural landscapes are more resistant to disturbances than population in pristine habitats or in pasturelands. This trend is however, not seen in populations with a high mobility.

Furthermore, on the life history traits aspect, populations with a lower mobility experienced a reduction of the age at first reproduction in disturbed habitats, while species with a higher mobility are experiencing a shorter generation time in habitats under higher human pressure. These results emphasize the importance of mobility in the response of animal population to human pressure.

**Table S8.** Summary of results of the residual method on demographic traits of animals that have a large migration range. Results are given in the following format: estimation of the effect (*P*-value). Underlined *P*-values are statistically significant (*i.e.* < 0.05) and italic P-values stand values between 0.05 and 0.10. *X* in the size column signified that no correlation was found between the demographic trait and size hence, there was no need for correction.

| Residual method for low mobility and high mobility animals | | | | |
|---|---|---|---|---|
| Trait | Human presence axis | Agricultural land-use | Size correction | Spatial correction |
| **Vital rates** | | | | |
| Survival rate SSD | -0.02 (0.35)<br>-0.01 (0.47) | -0.04 (0.46)<br>-0.02 (0.18) | YES<br>YES | YES<br>NO |
| Growth rate SSD | 0.02 (0.35)<br>0.01 (0.54) | 0.07 (0.26)<br>-0.02 (0.37) | YES<br>YES | YES<br>NO |
| Reproductive rate SSD | -0.05 (0.43)<br>0.01 (0.87) | 0.04 (0.30)<br>0.03 (0.71) | YES<br>NO | NO<br>NO |
| Survival rate of reproductive stages only SSD | 0.00 (0.91)<br>0.00 (0.93) | 0.00 (0.97)<br>-0.01 (0.51) | YES<br>YES | YES<br>NO |
| Growth rate of reproductive stages only SSD | 0.00 (0.98)<br>-0.01 (0.76) | -0.11 (0.33)<br>-0.05 (0.18) | NO<br>YES | NO<br>NO |
| **Life History Traits** | | | | |
| Reproductive rate | -0.02 (0.80)<br>0.09 (0.15) | 0.06 (0.74)<br>0.16 (0.23) | NO<br>NO | NO<br>YES |
| Degree of iteroparity | -0.08 (0.38)<br>-0.03 (0.57) | 0.03 (0.86)<br>-0.02 (0.88) | YES<br>NO | NO<br>YES |
| Generation time | 0.27 (0.22)<br>0.39 (0.02) | -0.09 (0.83)<br>0.79 (0.03) | YES<br>YES | YES<br>NO |
| Age at maturity | -0.05 (0.03)<br>-0.02 (0.71) | -0.17 (0.01)<br>-0.07 (0.50) | YES<br>YES | YES<br>YES |
| Survivorship | -0.02 (0.47)<br>-0.01 (0.40) | -0.02 (0.67)<br>0.00 (0.85) | NO<br>YES | YES<br>YES |

| | | | | |
|---|---|---|---|---|
| Probability of reproduction | 0.01 (0.17)<br>0.00 (0.84) | 0.00 (0.72)<br>0.02 (0.51) | YES<br>YES | YES<br>YES |
| Transient dynamics | | | | |
| Damping ratio | -0.01(0.64)<br>0.00 (0.86) | -0.03 (0.69)<br>-0.01 (0.87) | NO<br>YES | NO<br>YES |
| Period of oscillation | 0.00 (0.96)<br>0.00 (0.75) | -0.01 (0.92)<br>0.01 (0.71) | NO<br>YES | NO<br>YES |
| Reactivity | -0.01 (0.88)<br>0.01 (0.41) | 0.02 (0.88)<br>0.04 (*0.06*) | NO<br>NO | NO<br>NO |
| First step attenuation | -0.03 (*0.06*)<br>-0.01 (0.48) | -0.08 (<u>0.04</u>)<br>-0.02 (0.28) | NO<br>YES | NO<br>YES |

**Table S9.** Results of the MCMCglmm of the different demographic traits for plants against the two axes of HFP-PCA incorporating both phylogenetic corrections and correction for size. Results are given in the following format: estimation of the effect (*P*-value). Underlined *P*-values are statistically significant (*i.e.* < 0.05) and italic P-values stand values between 0.05 and 0.10. *X* in the size column signified that no correlation was found between the demographic trait and size hence, there was no need for correction.

| Phylogenetic correction for plants and animals | | | |
|---|---|---|---|
| Trait | Human presence axis | Agricultural land-use | Size |
| Vital rates | | | |
| Survival rate SSD | 0.00 (0.60)<br>-0.01 (0.22) | 0.00 (0.43)<br>-0.03 (0.16) | X<br>0.05 (0.42) |
| Growth rate SSD | 0.00 (0.60)<br>0.00 (0.93) | 0.00 (0.40)<br>-0.01 (0.60) | -0.01 (0.37)<br>0.00 (0.92) |
| Shrinkage rate SSD | 0.02 (0.26)<br>- | 0.00 (0.84)<br>- | -0.13 (<u>0.01</u>)<br>- |
| Reproductive rate SSD | 0.00 (0.90)<br>-0.02 (0.55) | 0.05 (0.25)<br>-0.01 (0.90) | 0.35 (<u>0.00</u>)<br>-0.03 (0.73) |
| Clonality rate SSD | 0.00 (0.66)<br>- | -0.01 (0.56)<br>- | X<br>- |
| Shrinkage rate of pre-reproductive stages SSD | 0.01 (<u>0.00</u>)<br>- | 0.02 (<u>0.03</u>)<br>- | X<br>- |
| Survival rate of reproductive stages only SSD | 0.00 (0.76)<br>0.00 (0.80) | 0.00 (0.69)<br>-0.01 (0.51) | X<br>0.05 (0.41) |
| Growth rate of reproductive stages only SSD | 0.01 (0.16)<br>0.00 (0.85) | 0.01 (0.18)<br>-0.06 (*0.09*) | -0.01 (0.44)<br>-0.02 (0.84) |
| Life History Traits | | | |
| Reproductive rate | 0.05 (0.23)<br>0.06 (0.25) | 0.07 (0.27)<br>0.13 (0.25) | 0.16 (*0.06*)<br>X |
| Degree of iteroparity | -0.05 (0.18)<br>0.00 (0.93) | -0.06 (0.31)<br>0.00 (0.99) | X<br>0.13 (0.15) |

| | | | |
|---|---|---|---|
| Generation time | -0.01 (0.45)<br>0.28 (*0.08*) | -0.01 (0.68)<br>0.50 (0.14) | 0.18 (<u>0.00</u>)<br>0.97 (<u>0.01</u>) |
| Age at first reproduction | -0.02 (0.26)<br>0.03 (0.38) | 0.01 (0.83)<br>0.05 (0.50) | 0.32 (<u>0.00</u>)<br>0.26 (<u>0.02</u>) |
| Survivorship | 0.00 (0.58)<br>-0.01 (0.37) | 0.01 (0.20)<br>-0.01 (0.81) | 0.04 (<u>0.05</u>)<br>0.02 (0.76) |
| Probability of reproduction | 0.00 (0.41)<br>0.00 (0.67) | 0.00 (0.49)<br>0.02 (0.43) | -0.03 (*0.08*)<br>X |
| Transient dynamics | | | |
| Damping ratio | 0.00 (0.50)<br>0.05 (0.15) | 0.01 (0.24)<br>0.08 (0.30) | -0.07 (<u>0.00</u>)<br>X |
| Period of oscillation | 0.02 (0.18)<br>0.00 (1.00) | 0.02 (0.34)<br>0.00 (0.99) | 0.23 (<u>0.00</u>)<br>0.00 (1.00) |
| Reactivity | -0.02 (0.34)<br>0.00 (0.92) | 0.00 (0.91)<br>0.03 (0.52) | 0.28 (<u>0.00</u>)<br>0.00 (0.99) |
| First step attenuation | 0.01 (*0.10*)<br>-0.01 (<u>0.02</u>) | 0.00 (0.95)<br>-0.04 (<u>0.01</u>) | 0.00 (0.77)<br>X |

**Table S10.** Results of the GAM of the different demographic traits for plants against the two axes of HFP-PCA incorporating correction for size and spatial autocorrelation. Results are given in the following format: estimation of the effect (*P*-value). Underlined *P*-values are statistically significant (*i.e.* < 0.05) and italic P-values stand values between 0.05 and 0.10. *X* in the size column signified that no correlation was found between the demographic trait and size hence, there was no need for correction. Rows with only X values meant that no spatial correction was detected and hence, that there was no need to correct for it.

| Spatial correction for plants and animals | | | |
|---|---|---|---|
| Trait | Human presence axis | Agricultural land-use | Size |
| Vital rates | | | |
| Survival rate SSD | -0.01 (0.00)<br>X | -0.03 (0.00)<br>X | X<br>X |
| Growth rate SSD | 0.00 (0.87)<br>X | -0.02 (0.01)<br>X | -0.02 (0.01)<br>X |
| Shrinkage rate SSD | 0.03 (*0.09*)<br>- | 0.02 (0.35)<br>- | 0.02 (0.50)<br>- |
| Reproductive rate SSD | 0.11 (0.00)<br>-0.14 (0.00) | 0.25 (0.00)<br>-0.58 (0.00) | 0.36 (0.00)<br>-0.09 (0.01) |
| Clonality rate SSD | 0.01 (*0.10*)<br>- | 0.02 (0.14)<br>- | X<br>- |
| Shrinkage rate of pre-reproductive stages SSD | 0.01 (0.16)<br>- | 0.01 (0.28)<br>- | X<br>- |
| Survival rate of reproductive stages only SSD | X<br>0.01 (0.27) | X<br>0.01 (0.59) | X<br>0.04 (0.00) |
| Growth rate of reproductive stages only SSD | 0.00 (0.67)<br>X | -0.01 (0.47)<br>X | -0.02 (0.02)<br>X |
| Life History Traits | | | |

| | | | |
|---|---|---|---|
| Reproductive rate | X<br>X | X<br>X | X<br>X |
| Degree of iteroparity | X<br>X | X<br>X | X<br>X |
| Generation time | -0.08 (0.00)<br>0.06 (0.78) | -0.08 (0.01)<br>-0.42 (0.45) | 0.06 (0.04)<br>0.80 (0.00) |
| Age at first reproduction | 0.00 (0.82)<br>-0.01 (0.66) | 0.01 (0.27)<br>-0.18 (0.15) | 0.01 (0.23)<br>0.18 (0.00) |
| Survivorship | -0.02 (0.01)<br>X | -0.01 (0.22)<br>X | 0.04 (0.00)<br>X |
| Probability of reproduction | X<br>0.02 (0.04) | X<br>0.13 (0.00) | X<br>X |
| Transient dynamics | | | |
| Damping ratio | 0.00 (0.87)<br>-0.07 (0.26) | 0.00 (0.97)<br>-0.01 (0.94) | -0.05 (0.00)<br>X |
| Period of oscillation | -0.03 (*0.08*)<br>0.00 (0.23) | 0.01 (0.76)<br>0.00 (0.75) | 0.16 (0.00)<br>0.00 (0.87) |
| Reactivity | 0.04 (0.13)<br>-0.04 (*0.09*) | 0.17 (0.00)<br>-0.24 (0.00) | 0.28 (0.00)<br>-0.04 (0.04) |
| First step attenuation | -0.01 (0.05)<br>X | -0.01 (*0.06*)<br>X | 0.00 (0.80)<br>X |

**Table S11.** Results of the MCMCglmm of the different demographic traits for animals with long migration range against the two axes of HFP-PCA incorporating both phylogenetic corrections and correction for size. Results are given in the following format: estimation of the effect (*P*-value). Underlined *P*-values are statistically significant (*i.e.* < 0.05) and italic P-values stand values between 0.05 and 0.10. *X* in the size column signified that no correlation was found between the demographic trait and size hence, there was no need for correction.

| Phylogenetic correction for low mobility and high mobility animals | | | |
|---|---|---|---|
| Trait | Human presence axis | Agricultural land-use | Size |
| Vital rates | | | |
| Survival rate SSD | -0.02 (0.41)<br>-0.01 (0.45) | -0.04 (0.47)<br>-0.02 (0.18) | 0.04 (0.84)<br>0.05 (0.55) |
| Growth rate SSD | 0.01 (0.40)<br>-0.01 (0.55) | 0.03 (0.63)<br>-0.02 (0.33) | X<br>-0.01 (0.90) |
| Reproductive rate SSD | -0.05 (0.42)<br>0.01 (0.87) | -0.04 (0.81)<br>0.03 (0.73) | -0.09 (0.73)<br>X |
| Survival rate of reproductive stages only SSD | 0.00 (0.89)<br>0.00 (0.92) | -0.02 (0.74)<br>-0.01 (0.51) | 0.06 (0.73)<br>0.05 (0.54) |
| Growth rate of reproductive stages only SSD | 0.00 (0.98)<br>-0.01 (0.75) | -0.11 (0.34)<br>-0.05 (0.19) | X<br>-0.02 (0.85) |
| Life History Traits | | | |
| Reproductive rate | -0.02 (0.80)<br>0.12 (*0.07*) | 0.06 (0.77)<br>0.22 (0.11) | X<br>X |
| Degree of iteroparity | -0.08 (0.40)<br>-0.04 (0.41) | 0.03 (0.89)<br>-0.04 (0.76) | 0.26 (0.26)<br>X |
| Generation time | 0.36 (0.33)<br>0.38 (<u>0.03</u>) | 0.39 (0.61)<br>0.79 (<u>0.04</u>) | 2.41 (<u>0.00</u>)<br>0.72 (<u>0.02</u>) |
| Age at first reproduction | -0.04 (*0.10*)<br>0.02 (0.67) | -0.10 (*0.09*)<br>0.03 (0.74) | 0.31 (0.28)<br>0.27 (<u>0.02</u>) |

| | | | |
|---|---|---|---|
| Survivorship | -0.02 (0.47)<br>-0.01 (0.44) | -0.04 (0.48)<br>0.00 (0.89) | X<br>0.02 (0.78) |
| Probability of reproduction | -0.01 (0.77)<br>0.01 (0.32) | -0.01 (0.93)<br>0.03 (0.20) | X<br>0.00 (0.97) |
| Transient dynamics | | | |
| Damping ratio | -0.01(0.66)<br>0.06 (0.19) | -0.03 (0.73)<br>0.11 (0.21) | X<br>-0.15 (0.29) |
| Period of oscillation | -0.01 (0.96)<br>0.00 (0.96) | -0.01 (0.96)<br>0.00 (0.92) | X<br>0.00 (0.97) |
| Reactivity | -0.01 (0.93)<br>0.01 (0.42) | 0.02 (0.87)<br>0.04 (*0.06*) | X<br>X |
| First step attenuation | -0.03 (0.04)<br>-0.01 (0.26) | -0.08 (0.03)<br>-0.03 (*0.10*) | X<br>0.01 (0.93) |
